# Characterization of SARS-CoV-2 Convalescent Patients’ Serological Repertoire Reveals High Prevalence of Iso–RBD Antibodies

**DOI:** 10.1101/2023.09.08.556349

**Authors:** Nicholas C. Curtis, Seungmin Shin, Andrew P. Hederman, Ruth I. Connor, Wendy F. Wieland-Alter, Steven Ionov, Jennifer Boylston, Josh Rose, Mrunal Sakharkar, Dana B. Dorman, John A. Dessaint, Lorraine L. Gwilt, Andrew R. Crowley, Jared Feldman, Blake M. Hauser, Aaron G. Schmidt, Alix Ashare, Laura M. Walker, Peter F. Wright, Margaret E. Ackerman, Jiwon Lee

## Abstract

While our understanding of SARS-CoV-2 pathogenesis and antibody responses following infection and vaccination has improved tremendously since the outbreak in 2019, the sequence identities and relative abundances of the individual constituent antibody molecules in circulation remain understudied. Using Ig-Seq, we proteomically profiled the serological repertoire specific to the whole ectodomain of SARS-CoV-2 prefusion-stabilized spike (S) as well as to the receptor binding domain (RBD) over a 6-month period in four subjects following SARS-CoV-2 infection before SARS-CoV-2 vaccines were available. In each individual, we identified between 59 and 167 unique IgG clonotypes in serum. To our surprise, we discovered that ∼50% of serum IgG specific for RBD did not recognize prefusion-stabilized S (referred to as iso–RBD antibodies), suggesting that a significant fraction of serum IgG targets epitopes on RBD inaccessible on the prefusion-stabilized conformation of S. On the other hand, the abundance of iso–RBD antibodies in nine individuals who received mRNA-based COVID-19 vaccines encoding prefusion-stabilized S was significantly lower (∼8%). We expressed a panel of 12 monoclonal antibodies (mAbs) that were abundantly present in serum from two SARS-CoV-2 infected individuals, and their binding specificities to prefusion-stabilized S and RBD were all in agreement with the binding specificities assigned based on the proteomics data, including 1 iso–RBD mAb which bound to RBD but not to prefusion-stabilized S. 2 of 12 mAbs demonstrated neutralizing activity, while other mAbs were non-neutralizing. 11 of 12 mAbs also bound to S (B.1.351), but only 1 maintained binding to S (B.1.1.529). This particular mAb binding to S (B.1.1.529) 1) represented an antibody lineage that comprised 43% of the individual’s total S-reactive serum IgG binding titer 6 months post-infection, 2) bound to the S from a related human coronavirus, HKU1, and 3) had a high somatic hypermutation level (10.9%), suggesting that this antibody lineage likely had been elicited previously by pre-pandemic coronavirus and was re-activated following the SARS-CoV-2 infection. All 12 mAbs demonstrated their ability to engage in Fc-mediated effector function activities. Collectively, our study provides a quantitative overview of the serological repertoire following SARS-CoV-2 infection and the significant contribution of iso–RBD antibodies, demonstrating how vaccination strategies involving prefusion-stabilized S may have reduced the elicitation of iso–RBD serum antibodies which are unlikely to contribute to protection.

## Introduction

Since its emergence three years ago, severe acute respiratory syndrome coronavirus 2 (SARS-CoV-2) has caused nearly 7 million deaths and 770 million illnesses globally. SARS-CoV-2 infects its target cells by fusing to cell membranes through the receptor binding domain (RBD) on trimeric spike (S) protein that contacts the host cell receptor, angiotensin-converting enzyme 2 (ACE2) (1, 2). After membrane fusion and proteolytic cleavage, the S1 subunit, which includes the RBD, dissociates (3, 4), which has been shown to expose novel epitopes on RBD not accessible in the prefusion conformation (5). The majority of approved vaccines is based on prefusion-stabilized S, including BNT162b2 (Pfizer-BioNTech) (6), mRNA-1273 (Moderna) (7), and Ad26.CoV2.S (Janssen) (8), due to its greater stability and potentially increased immunogenicity (9). Current trials are also exploring using RBD alone as an immunogen (10–15) to focus immune responses to RBD, where neutralizing antibodies predominantly bind (16–18).

Over the past few years, a number of studies have elucidated how the antibody repertoire to SARS-CoV-2 is generated. Following infection, memory B cells specific to SARS-CoV-2 S accumulate and undergo further rounds of somatic hypermutation (SHM) (19, 20), which significantly increases between 1- and 6-months post-infection (21) and reach similar levels to those seen in B cells specific to other antigens such as those following influenza vaccination (22) and respiratory syncytial virus infection (23). These SARS-CoV-2 S-reactive memory B cells predominantly target the S2 subunit, with a minority targeting RBD and N-terminal domain (NTD) (24). Circulating immunoglobulins in serum also predominantly target outside of the RBD (both S2 and NTD) (25, 26) and peak in abundance roughly 16-30 days following infection (27). While SARS-CoV-2 S- and RBD-reactive memory B cells encoding antibodies with neutralization activity continue to persist many months following infection, serum antibody binding and neutralization titers wane rapidly (20), and in some individuals revert to pre-infection levels of neutralization (19). The rates of decay are more rapid for S-reactive serum IgM and IgA (27, 28); in most individuals neutralizing IgG responses can be detected over a year after infection (29, 30).

Recent studies have focused on the impact of pre-established antibody repertoire generated prior to the onset of the pandemic. SARS-CoV-2 is a member of the *Betacoronavirus* genus, which also includes other seasonal human coronaviruses (hCoVs), HKU1 and OC43. Studies have shown that antibody responses elicited in approximately 20% of SARS-CoV-2 infected individuals, predominantly those with severe symptoms (31, 32), show varying degrees of cross-reactivity to hCoVs. Another study has shown that the early memory B cell response following SARS-CoV-2 infection derives from hCoV-reactive B cells (20). These hCoV-cross-reactive antibodies may have been elicited by exposures to hCoVs before the SARS-CoV-2 pandemic (33). S2 is the target of many of these antibodies and is more conserved among *Betacoronavirus*. It has been demonstrated that SARS-CoV-2 naïve individuals (particularly children (34), who often have more recently been exposed to hCoV (33)) have memory B cells recognizing SARS-CoV-2 S, which is further boosted by SARS-CoV-2 infection (34, 35). Similar observations have been made from circulating antibodies, with S2-reactive antibodies in serum being back-boosted (36), presumably from earlier hCoV infection. These hCoV cross-reactive antibodies lack neutralization activity (37–39), and the presence of hCoV cross-reactive antibodies prior to SARS-CoV-2 infection did not lead to differences in serum titers following infection or frequency of infection in SARS-CoV-2 naïve individuals, implying their limited role in protection (31).

While the SARS-CoV-2 pandemic is mitigated by available vaccines and therapeutics, concerns still remain due to the continued rise of variants of concern (VoCs) with increased infectivity (40–47) and which contain mutations on antibody binding residues, particularly in the RBD (40, 47–60), to evade antibody recognition. Many of these variants become more transmissible (45, 61–63) with key mutations, such as D614G, characteristic of every variant after the B.1.1.7 variant (41, 43, 45, 56, 61), and E484K (44, 51) found in B.1.351 and B.1.1.529. Consequently, studies have shown that antibodies from previously infected or vaccinated individuals have reduced binding and decreased neutralization against the B.1.1.7 (53, 64), B.1.351 (53, 64), B.1.617.2 (65–67), and B.1.1.529 variants (68, 69). To address the reduced neutralization activity against circulating strains, there are ongoing efforts to update vaccines as the virus mutates by either immunizing with a contemporary SARS-CoV-2 variant (70) or multivalent vaccines (71, 72), or by developing vaccination strategies to elicit higher titers of cross-reactive antibodies by immunizing with only the S1 (73), S2 (74), RBD (75), or a mixture of immunogenic SARS-CoV-2 peptides (76).

Using Ig-Seq, a proteomics-based serological antibody repertoire identification and quantitation technology (77–80), we profiled the secreted antibody molecules specific to prefusion-stabilized S and RBD in four unvaccinated, SARS-CoV-2 convalescent individuals with varying severity of illness, that have been previously studied (38, 81–83). Interestingly, we found that ∼50% of serum antibodies affinity-purified with RBD cannot bind to prefusion-stabilized S (‘iso–RBD’ antibodies). In contrast, vaccinated individuals had substantially lower levels of these iso–RBD antibodies. By recombinantly expressing representative antibodies from abundant serum antibody clonotypes in our Ig-Seq data, we identified one antibody clonotype from a donor that constituted as much as 43% of the serum IgG binding titer to S; this clonotype had a very high level of SHM and also bound strongly to HKU1 S, suggesting that this antibody lineage likely have been elicited previously by pre-pandemic coronavirus before being re-activated by the SARS-CoV-2 infection.

## Results

### Features of the Anti-SARS-CoV-2 S Serological IgG Repertoires

Serum samples from four individuals were collected at 1-month (M1), 3-months (M3), and 6-months (M6) following SARS-CoV-2 infection for the Ig-Seq analysis (**Fig. 1A** and **Supplementary Table 1**) before SARS-CoV-2 vaccines were available. From each serum sample, SARS-CoV-2 S-reactive serum IgG was purified using two separate affinity chromatography columns; one with the whole ectodomain of prefusion-stabilized S (HexaPro) (84) immobilized and the other one with the RBD-SD1 (referred to as ‘RBD’) (85) immobilized, both from the Wuhan-Hu-1 isolate (Wu), which is likely the strain infecting the donors based on their infection dates. The inclusion of both antigens in the experimental design was motivated by the facile identification of regions targeted by individual serum antibody clonotypes (*i.e.*, if the same antibody clonotypes are detected in the eluates of both prefusion-stabilized S and RBD pulldowns, that would indicate that the clonotype targets the RBD region on S). Eluates from each pulldown were analyzed separately by high-resolution liquid chromatography-tandem mass spectrometry (LC-MS/MS). MS spectra generated from each subject’s samples were annotated using the matched donor-specific database of putative B cell receptor sequences generated by Next-Gen Sequencing of peripheral B cells isolated at M1 following infection (BCR-Seq) (**Supplementary Table 2**).

**Figure 1.**
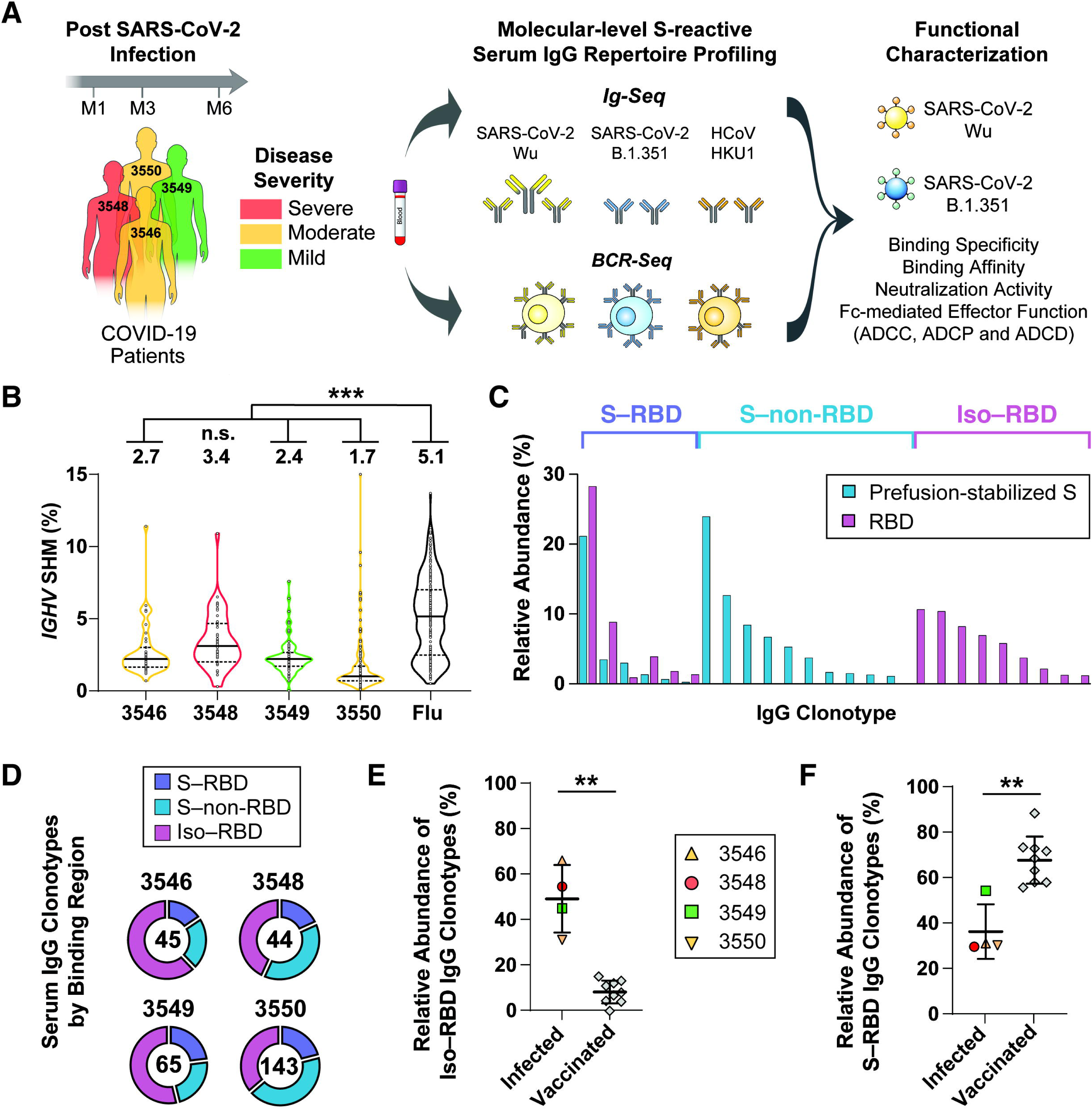
Profiling the serum IgG repertoire to SARS-CoV-2 S. **A.** Experimental design. Blood was collected from four donors with either severe, moderate, or mild symptoms 1-month (M1), 3-months (M3), and 6-months (M6) following SARS-CoV-2 infection. Combining Ig-Seq with paired-chain single B cell sequencing, BCR-Seq, serum antibody clonotypes were proteomically identified following affinity purification with prefusion-stabilized S or RBD. Selected abundantly present serum antibodies were recombinantly expressed and tested for binding affinity and specificity, neutralization activity, and Fc-mediated effector functions against SARS-CoV-2 Wu and B.1.351. **B.** *IGHV* SHM levels of the anti-SARS-CoV-2 S serological IgG repertoires. The influenza data set is based on HA-reactive serum antibodies identified in four adults following trivalent influenza vaccination (79). Black horizontal lines represent medians (solid) and quartiles (dotted), with mean values included above each plot. Statistical significance is determined by Dunn’s multiple comparisons test (*** *p* < 0.001). **C.** Representative serological repertoire from subject-3548 at M1. Each bar indicates individual IgG clonotypes, and its height represents relative abundance. **D.** IgG clonotypic diversity based on different binding regions at M1. The total number of clonotypes identified is included at the center of each plot. **E-F.** Relative abundance of iso–RBD serum IgG in the eluates from RBD pulldowns (**E**) and relative abundance of S–RBD serum IgG in the eluates from prefusion-stabilized S pulldowns (**F**) at M1. Nine vaccinated individuals are included as a comparison. Relative abundance is calculated using the proteomics-based quantification of each antibody clonotype in serum. Mean values are represented as horizontal lines with the error bars indicating SD. Statistical significance is determined by a Mann-Whitney U test. (** *p* < 0.01).

Inspection of the serological repertoires for all of the donors revealed several insights. First, the number of unique serum IgG clonotypes identified in the eluates of prefusion-stabilized S pulldowns ranged between 26 and 103, and between 38 and 95 serum IgG clonotypes were also identified in the eluates of RBD pulldowns (the total number of unique clonotypes ranged between 59 and 167) (**Supplementary Fig. 1** and **Supplementary Data**). This level of clonotypic diversity of the serum antibody repertoire is similar to what has been observed in SARS-CoV-2 (26, 86) and influenza (79, 80, 87, 88). Additionally, the serological repertoire to S of these convalescent donors was polarized, with the top 16% of the most abundant clonotypes accounting for more than 60% of the total abundance in the entire S-reactive IgG repertoire (**Supplementary Table 3**). Next, the average *IGHV* SHM levels for S-reactive serum antibodies ranged from 1.7% to 2.7% for three (subject-3546, subject-3549, and subject-3550) of the four subjects (**Fig. 1B**), similar to what we have previously observed in infants receiving flu vaccines (88). For subject-3548, who had the most severe symptoms, the SHM level (3.4%) approached those seen in adults following influenza vaccination (79). Correlation between increased SHM levels with increased disease severity has been previously reported in the B cell repertoire, explained by the persistence of viral antigens for prolonged germinal center reactions (20, 89, 90), but this is the first observation in the context of IgG in circulation. Other features including CDRH3 length, charge, and hydrophobicity, and *IGHV*-*IGHJ* usage were similar among the infected individuals (**Supplementary Fig. 2** and **Supplementary Data**).

### Prevalence of Iso–RBD Serum Antibodies

Binding regions targeted by individual serum IgG clonotypes were assigned based on the proteomics data and designated as 1) ‘S–RBD’, if detected in the eluates of both the prefusion-stabilized S and RBD pulldowns (*i.e.*, bind RBD on prefusion-stabilized S) 2) ‘S–non-RBD’, if detected only in the eluates from the prefusion-stabilized S pulldown (*i.e.*, bind non-RBD regions on prefusion-stabilized S), and 3) iso–RBD, if detected only in the eluates from the RBD pulldown (*i.e.*, bind regions on RBD not accessible on prefusion-stabilized S) (**Fig. 1C**). When we examined the binding region distribution of the serum antibody clonotypes at M1, we noticed that an unexpectedly high number of serum IgG clonotypes were categorized as iso–RBD across the four subjects (on average 49±11%), whereas S–RBD and S–non-RBD clonotypes comprised 17±3% and 34±10%, respectively (**Fig. 1D**). This observation was surprising to us because such high prevalence of iso–RBD antibodies in sera has not been reported before, although a couple of studies have shown examples of monoclonal antibodies (mAbs) identified through phage panning from synthetic and alpaca-derived antibody libraries that bind to recombinantly expressed soluble RBD but not to prefusion-stabilized S (91, 92). We considered the possibility that these iso–RBD antibody clonotypes may be low in abundance and were not detected in the eluate of prefusion-stabilized S pulldown due to limitations in the detection sensitivity of Ig-Seq. However, many of the iso–RBD clonotypes in each donor were highly abundant based on the quantitation through the proteomics analysis, with the most abundant iso–RBD clonotype (clonotype-81735) comprising nearly 16% of the total IgG antibodies binding to RBD in subject-3549 (**Supplementary Table 4**). When we summed the relative abundance of all iso–RBD clonotypes, they accounted for nearly 50% (ranging between 31% and 66%) of serum antibodies targeting RBD (**Fig. 1E**). This finding suggests that nearly half of all RBD-directed serum antibodies elicited by the infection target epitopes not accessible in the prefusion-stabilized conformation of S.

As one mechanism for the elicitation of iso–RBD antibodies may be in recognition of novel epitopes on RBD exposed upon S1 detachment (5), we hypothesized that iso–RBD antibodies would be rare in infection-naïve individuals who have received COVID-19 vaccines based on prefusion-stabilized S immunogens. In a separate study, we profiled the S-reactive serum IgG from nine infection-naïve individuals with Cystic Fibrosis four weeks after the second dose from a two-dose vaccination regime (**Supplementary Table 5**). In these subjects, we observed significantly lower abundance of iso–RBD antibodies, which comprised only 8±5% of the serological repertoire (**Fig. 1E**). In the four convalescent individuals, S–RBD clonotypes accounted for 36% of the antibodies by abundance (ranging between 30% and 54%) (**Fig. 1F**), consistent with expectations based on observations from others (18, 26, 93, 94). Conversely, the vaccinated donors had significantly higher abundance of S–RBD antibodies, averaging 68±10%, explained by the decreased abundance of iso–RBD antibodies (**Fig. 1F**).

Next, we purified RBD-reactive IgA from serum at M1 from all four convalescent donors using the RBD-immobilized affinity column (**Supplementary Fig. 3A**). The anti-RBD serum IgA repertoire had fewer unique clonotypes compared to the IgG repertoire, ranging between 11 and 24 clonotypes (**Supplementary Data**). When we examined individual IgG and IgA clonotypes across the four donors, we noted that most antibody clonotypes were found in circulation only as IgG (designated as ‘IgG-only’) or IgA (designated as ‘IgA-only’), which comprised 77±11% and 17±8% of all clonotypes, respectively (**Supplementary Fig. 3B**). Only a small number of antibody clonotypes were detected in circulation as IgG as well as IgA (designated as ‘IgG+IgA’), ranging between 2% and 9% (on average, 6%), illustrating the distinctive nature of serum IgG and IgA repertoires. We noted that IgA-only clonotypes had higher SHM levels than IgG-only clonotypes (**Supplementary Fig. 3C**). This observation of the secreted IgA antibody clonotypes being more mature aligns with previous observations of S-reactive IgA B cells having higher SHM levels compared to the IgG B cells (95). We also observed abundance of iso–RBD IgA clonotypes as we did not see any significant differences between the binding regions of IgG-only and IgG+IgA clonotypes (**Supplementary Fig. 3D**).

### Dynamics of the SARS-CoV-2 S-Reactive Serum IgG Repertoire

As expected, the individuals with more severe symptoms showed higher serum IgG binding titers to S and RBD (**Fig. 2**, **A** and **B**). Over the course of 6 months, titers to S and RBD decayed by 58±11% and 66±12%, respectively, across the four donors, consistent with observations reported by others (27, 29, 60, 96–100). Serum neutralization titers measured with pseudotyped vesicular stomatitis virus (VSV) bearing the SARS-CoV-2 S also decreased by 92±6%, on average, over 6 months (**Fig. 2C**). We observed a strong positive correlation (Spearman’s ρ = 0.861) between the serum IgG binding titer to RBD and the neutralization activity measured at each time point (**Fig. 2D**), consistent with the observations of RBD-targeting antibodies contributing to the serum neutralization activity (101–105). Next, to quantitate serum IgG’s contribution to the total serum neutralization activity, we depleted all non-IgG isotypes and observed a 35-fold reduction in neutralizing activity against pseudotyped VSV-SARS-CoV-2 at M1, a 10-fold reduction at M3, and a 5-fold reduction at M6 (**Fig. 2E**) suggesting that non-IgG serum antibody isotypes contribute substantially to the SARS-CoV-2 neutralizing response (106, 107), especially during early time points.

**Figure 2.**
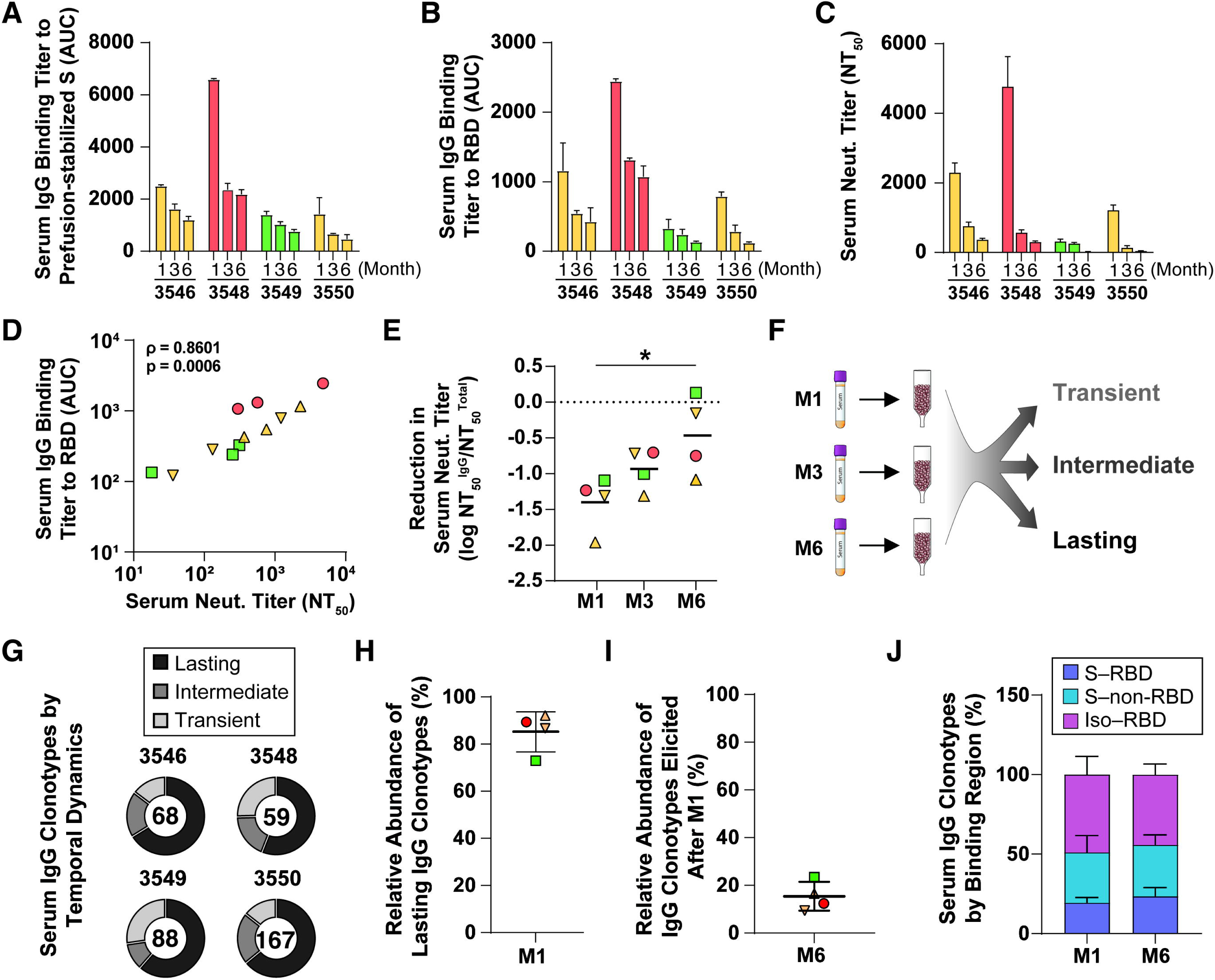
Longitudinal analysis of the anti-SARS-CoV-2 S serological IgG repertoires. **A.** Serum IgG binding titers to prefusion-stabilized S measured across all samples. **B.** Serum IgG binding titers to RBD measured across all samples. **C.** Serum neutralizing titers against pseudotyped VSV SARS-CoV-2 measured across all samples. **D.** Correlation between neutralization activity and serum binding titer to RBD. A two-sided parametric Spearman’s correlation test was used for statistical analyses. **E.** Reduction of neutralizing activity after depleting all antibody isotypes except IgG. Black horizontal lines represent means, and the dashed line indicates no reduction. Statistical significance is determined by a Mann-Whitney U test. (* *p* < 0.05). **F.** Definition of lasting, intermediate, and transient clonotypes used in this study. **G.** IgG clonotypic diversity based on different temporal dynamics. The total number of clonotypes is included at the center of each plot. **H.** Relative abundance of lasting clonotypes detected in the eluates from M1 prefusion-stabilized S pulldown. **I.** Relative abundance of antibody clonotypes in the eluates from M6 prefusion-stabilized S pulldown that were first detected after M1. **J.** Breakdown of IgG clonotypes detected in M1 and M6 by their binding regions. The error bars indicate SD. For **A**, **B**, and **C** all samples run in technical duplicate, and the error bars indicate SD. For **H**, and **I**, relative abundance is calculated using the proteomics-based quantification of each antibody clonotype in serum. Mean values are represented as horizontal lines with the error bars indicating SD.

Next, we designated IgG clonotypes as lasting (detected at M6), intermediate (detected at M3 but not at M6), or transient (detected only at M1) (**Fig. 2F**). At the clonotypic diversity level, lasting antibody clonotypes comprised between 56% and 66% (on average, 62%) (**Fig. 2G**), and the total abundance of all lasting clonotypes detected at M1 constituted 85±9% of the IgG response to S (**Fig. 2H**); in other words, most serum antibodies detected at M1 lasted in circulation over 6 months. When we quantified the total abundance of new antibody clonotypes elicited after M1 that were still present at M6, they constituted between 9% and 23% (on average, 15%) of the IgG response (**Fig. 2I**), indicating the composition of the serological repertoires at M6 is highly similar to those at M1, which highlight the relatively constant nature of the serological repertoire during the 6 months following infection. The clonotypic diversity remained steady between M1 and M6, leading to the binding regions also staying relatively unchanged (**Fig. 2J**).

### Features of Serum Antibodies Cross-Reactive to SARS-CoV-2 B.1.351 and HCoV HKU1

We characterized the serum IgG response in our four donors against the RBD (B.1.351), which contains three mutations (K417N, E484K, and N501Y). We observed lower serum IgG binding titers against the RBD (B.1.351) (**Fig. 3A**), and neutralizing activity was lower by a factor of 10.2 on average against pseudotyped VSV-SARS-CoV-2 (B.1.351) (**Fig. 3B**). Next, we affinity-purified RBD (B.1.351)-reactive IgG from M1 serum from each of the four convalescent donors using the RBD (B.1.351)-immobilized affinity column and analyzed via Ig-Seq to determine which IgG clonotypes are cross-reactive to B.1.351. In the serum IgG repertoire, between 24% and 44% (on average, 33%) of the antibody clonotypes from the RBD-pulldown were cross-reactive (**Fig. 3C**). When we accounted for the relative abundance of each clonotype, we found that between 36% and 85% (on average, 55%) of the serum IgG elicited by the infection also recognized RBD (B.1.351) (designated as ‘Wu+B.1.351 CR’) (**Fig. 3D**). This abundance of Wu+B.1.351 CR antibodies on the basis of the proteomic analysis was strongly correlated (*p* < 0.001) with the serum IgG binding titers to RBD (B.1.351) (**Fig. 3E**). A larger fraction of S–RBD clonotypes (between 50% and 75%; on average, 66%) were Wu+B.1.351 CR whereas the majority of iso–RBD clonotypes were specific to RBD (Wu) (designated as ‘Wu-specific’) (between 14% and 34%; on average, 23%) (**Fig. 3F**), implying iso–RBD antibodies target epitopes that are mutated on RBD (B.1.351). We note that the K417 residue, which is mutated to N417 in the B.1.351 variant, is located in one of the regions that becomes increasingly accessible on RBD when S1 detaches (5).

**Figure 3.**
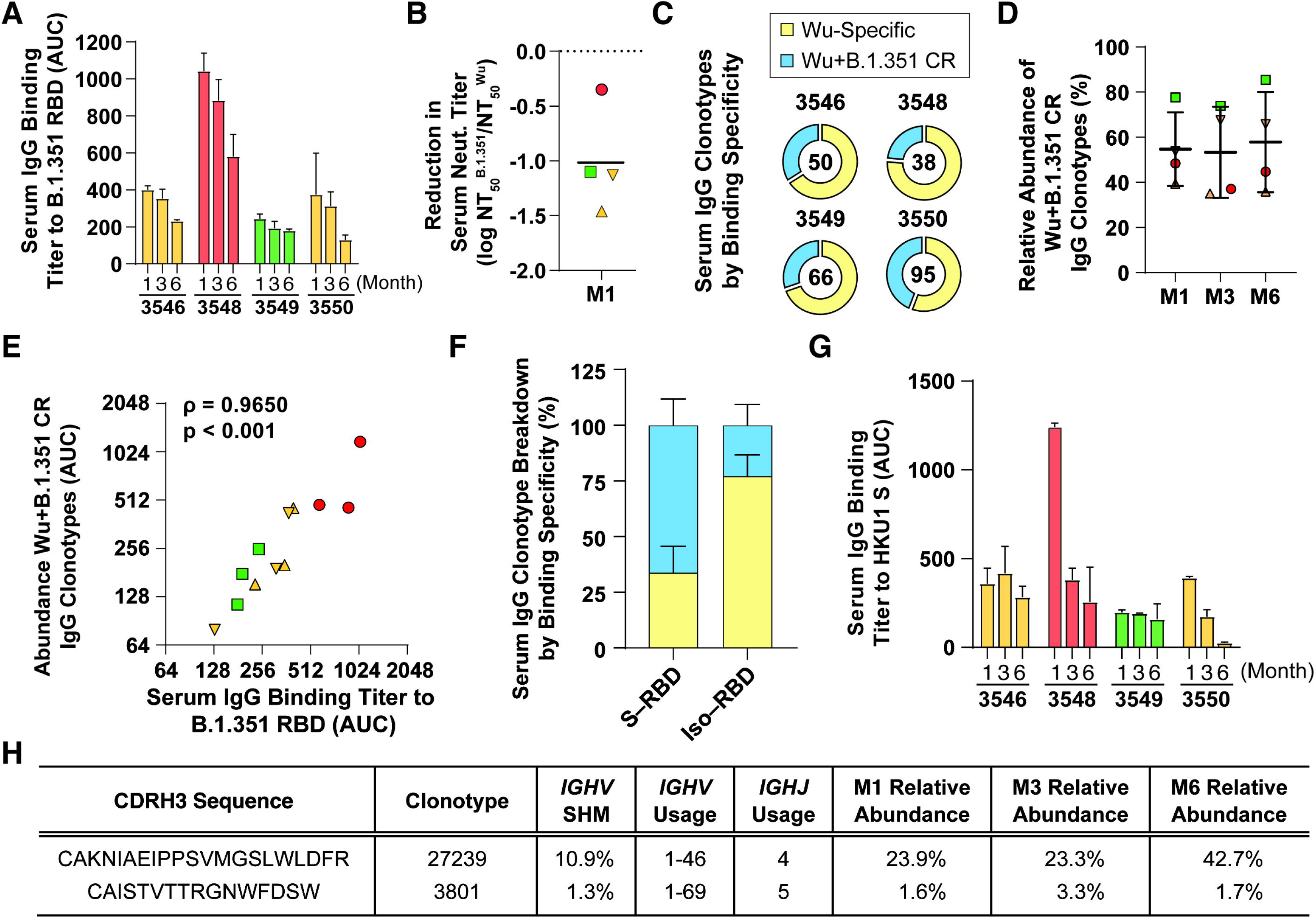
Cross-reactivity of RBD-reactive serum IgG repertoire. **A.** Serum IgG binding titer against RBD (B.1.351). **B.** Comparison of serum neutralizing titers against VSV-SARS-CoV-2 Wu and B.1.351 at M1. Black horizontal line represents the mean, and the dashed line indicates no reduction. **C.** Frequency of Wu-specific and Wu+B.1.351 CR clonotypes in the RBD-pulldown. The total number of clonotypes is included at the center of each plot. **D.** Relative abundance of Wu+B.1.351 CR clonotypes in the RBD-pulldown. Relative abundance is calculated using the proteomics-based quantification of each antibody clonotype in serum. Mean values are represented as horizontal lines with the error bars indicating SD. **E.** Correlation between serum IgG binding titers to RBD (Wu) scaled by abundance of Wu+B.1.351 CR antibodies based on the proteomics data and serum IgG binding titers to RBD (B.1.351). **F.** Frequency of Wu-specific and Wu+B.1.351 CR antibody clonotypes that are S–RBD or iso– RBD. The error bars indicate SD. **G.** Serum IgG binding titer against HKU1 S. **H.** Summary of HKU1 S cross-reactive antibodies. For **A** and **G**, all samples run in technical duplicate, and the error bars indicate SD.

Subsequently, we characterized serum antibody cross-reactivity to the S from HKU1, which shares 29% identity (40% similarity) with SARS-CoV-2 S. There is a higher degree of homology between these proteins in the S2 subunit where some regions have over 75% similarity (108–110). First, measured serum IgG binding titers to HKU1 S showed only subject-3548 had a robust level of serum antibodies that reacted to HKU1 S (**Fig. 3G**). From subject-3548, we affinity-purified serum IgG using HKU1 S and proteomically analyzed the eluates. We identified two clonotypes in the eluates of both HKU1 S and SARS-CoV-2 S pulldowns (**Fig. 3H**). One was the most abundant antibody clonotype from subject-3548 (clonotype-27239), accounting for 23.9% of the total serum IgG reactive to SARS-CoV-2 at M1. The relative abundance of this clonotype in circulation increased at M6 (up to 42.7%), showing a delayed rate of decay in comparison to other antibody clonotypes. This clonotype used *IGHV* 1-46 and *IGHJ* 4, with a very high level of SHM (10.9%), which is a feature of a recall response; this antibody lineage likely was elicited previously by pre-pandemic coronavirus and was re-activated following the SARS-CoV-2 infection. The other clonotype used *IGHV* 1-69 and *IGHJ* 5, had a SHM level of around 1.3% and comprised a small fraction (between 1.6% and 3.3%) of the total serum IgG reactive to SARS-CoV-2 S.

### Functional Characteristics of Representative Serum mAbs

We recombinantly expressed a representative mAb from each of the 12 abundant serum IgG clonotypes (including the most abundant ones): 9 S–non-RBD, 2 S–RBD, and 1 iso–RBD mAbs (**Table 1**) from two donors with more severe symptoms (subject-3546 and subject-3548). When tested via ELISA, all 12 mAbs bound to prefusion-stabilized S and RBD in perfect agreement with the binding regions designated based on the proteomics data; 2 S–RBD mAbs bound both prefusion-stabilized S and RBD, 9 S–non-RBD mAbs bound only to prefusion-stabilized S, and 1 iso–RBD mAb bound only to RBD with EC_50_ values ranging from ∼0.1 to 130 nM (**Fig. 4A**). The iso–RBD mAb, 3548-RBD-15, also bound to S1 while not binding to 3 different recombinantly expressed prefusion-stabilized S constructs as expected (**Supplementary Table 6**). 3548-RBD-15 also did not bind to RBD (B.1.351), which aligns with our earlier observation that iso–RBD antibodies are likely Wu-specific.

**Table 1.**
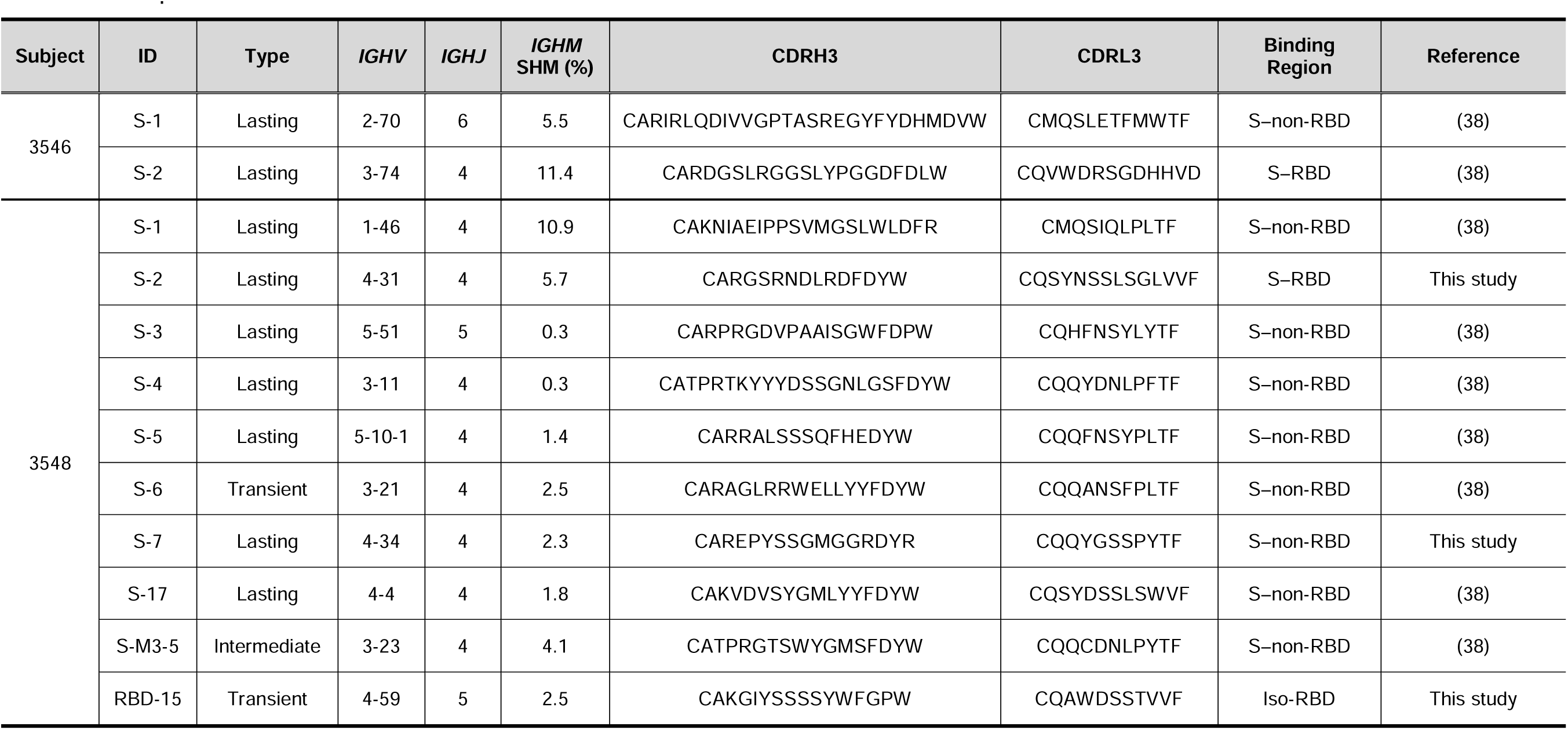
Summary of representative mAbs. mAb ID indicates reactivity (S or RBD) and rank of abundance in each donor’s M1 serum, except for 3548-S-M3-5 which is based on M3 serum.

**Figure 4.**
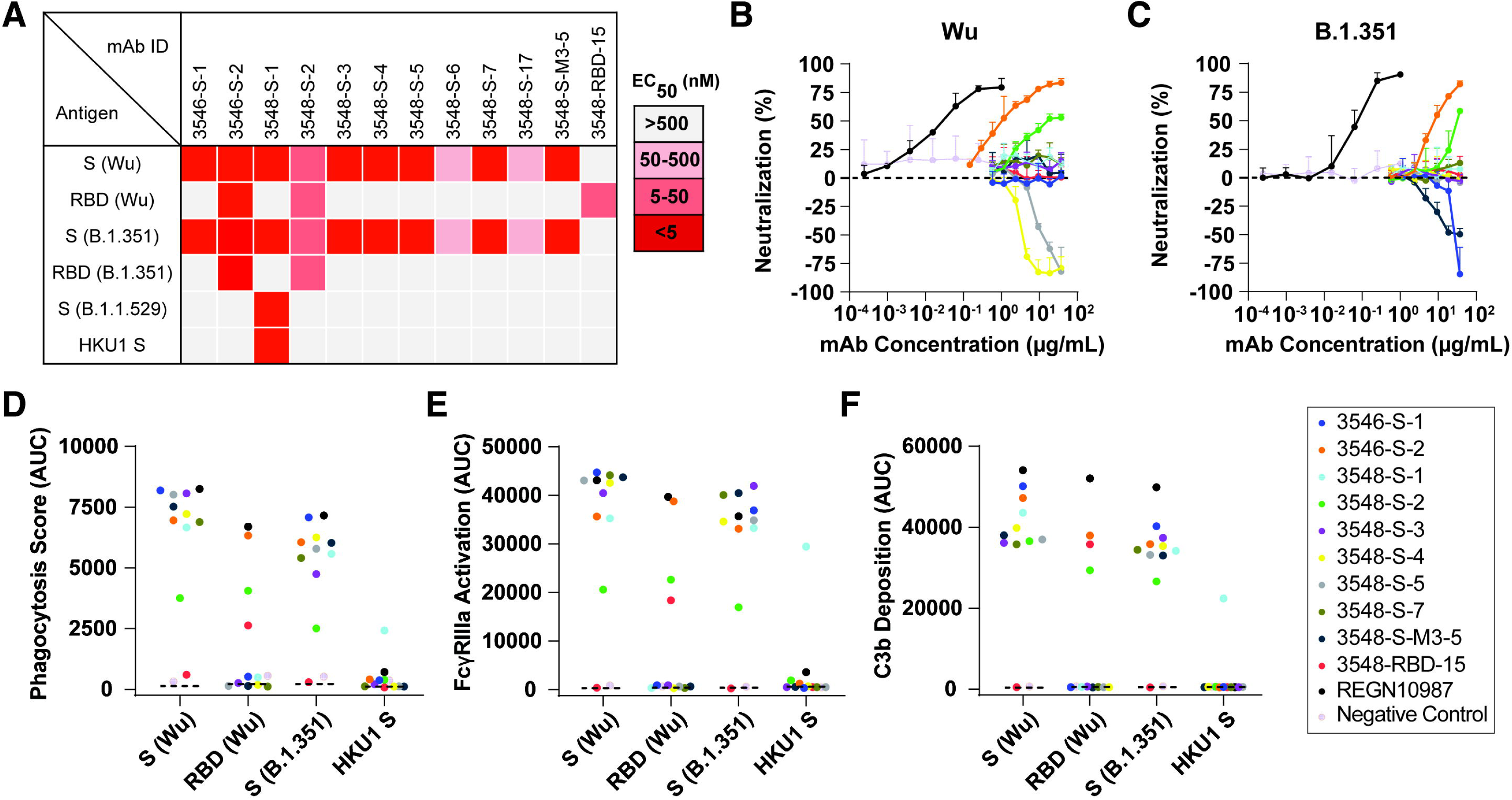
Biochemical and functional characterization of SARS-CoV-2 S- or RBD mAbs. **A.** Binding specificity and affinity measured by ELISA. EC_50_ values for each mAb to prefusion-stabilized S (Wu), RBD (Wu), prefusion-stabilized S (B.1.351), RBD (B.1.351), prefusion-stabilized S (B.1.1.529), and HKU1 S are shown. All assays were performed in triplicate. **B-C.** Neutralization activity of mAbs against pseudotyped VSV SARS-CoV-2 (Wu) (**B**) and VSV-SARS-CoV-2 (B.1.351) (**C**). CR3022, a non-neutralizing mAb was used as a negative control. The error bars indicate SD. **D-F.** Antibody-dependent cellular phagocytosis (ADCP) (**D**), FcγRIIIa activation (ADCC) (**E**), and antibody-dependent complement deposition (ADCD) (**F**) activity of mAbs against SARS-CoV-2 prefusion-stabilized S (Wu), RBD (Wu), prefusion-stabilized S (B.1351), and HKU1 S. REGN10987 IgG2σ (127) was used as a null Fc-mediated effector function control. The dotted lines indicate the no antibody control.

The other 11 mAbs showed cross-reactivity to S (B.1.351). 3546-S-2 and 3548-S-2 also showed cross-reactivity to RBD (B.1.351), which was expected as those two clonotypes were also detected in the eluate of the RBD (B.1.351) pulldown (**Supplementary Data**). Against prefusion-stabilized S (B.1.1.529), only 3548-S-1 maintained binding. 3548-S-1 is the representative mAb from clonotype-27239, which was also detected in the eluate of HKU S pulldown. As expected, 3548-S-1 also bound to HKU1 S, further validating our classification of binding specificities based on the proteomics data. We selected 10 mAbs with a binding EC_50_ below 50 nM for their binding kinetics (**Supplementary Table 7** and **Supplementary Fig. 4**). K_D_ values for these mAbs ranged between 0.087 nM to 63 nM, in alignment with our ELISA data. For mAbs with cross-reactivity to B.1.351 or HKU1, K_D_ values were similar between antigens from different variants.

Despite their strong binding affinities, only 2 S–RBD mAbs, 3546-S-2 and 3548-S-2, showed neutralization activity against pseudotyped VSV-SARS-CoV-2 (Wu) (**Fig. 4B**) and VSV-SARS-CoV-2 (B.1.351) (**Fig. 4C**). Given the majority of these abundantly present serum mAbs are lacking neutralization activity, other serum antibodies low in abundance or in different isotypes likely contribute to the observed serum neutralization titers (**Fig. 2C**). More importantly, 4 S– non-RBD mAbs, 3546-S-1, 3548-S-4, 3548-S-5, and 3548-S-M3-5, actually enhanced the infectivity of VSV-SARS-CoV-2 (Wu) or VSV-SARS-CoV-2 (B.1.351) pseudoviruses. It has been previously reported that some mAbs targeting NTD can improve the interaction between S and ACE2, which results in enhanced infectivity measured in neutralization assays (111, 112). Our observation is the first time such enhancing mAbs have been isolated directly from serum (*i.e*., functionally secreted antibody).

As non-neutralizing antibodies may still confer protection through Fc-mediated immune functions, we measured the antibody-dependent cellular phagocytosis (ADCP), cellular cytotoxicity (ADCC), and complement deposition (ADCD) activities of these mAbs (**Fig. 4**, **D-F** and **Supplementary Fig. 5**). All 10 mAbs showed their ability to engage in effector functions in various capacities, with 3548-S-2 and 3548-RBD-15 showing the weakest Fc-mediated effector functions; these 2 mAbs also had weaker binding affinities. Cross-reactive mAbs showed similar levels of ADCP, ADCC, and ADCD against Wu and B.1.351. The HKU1 S cross-reactive mAb, 3548-S-1, also showed robust ADCP, ADCC, and ADCD activities against HKU1. The ADCC activity of mAbs, which bind to SARS-CoV-2 S, correlated with the K_D_ from BLI, whereas ADCP and ADCD activities did not significantly correlate (**Supplementary Fig. 6**). Together, these results indicate that abundant SARS-CoV-2 antibodies in infected donors are mostly non-neutralizing but may still be able to elicit effective Fc-mediated functions in physiologically relevant settings.

## Discussion

We used proteomics-based quantitative analysis of serum antibodies (Ig-Seq) combined with high-throughput sequencing of B cell transcripts (BCR-Seq) to characterize the serological repertoire to SARS-CoV-2 S following infection in four individuals. Although the generalizability of our findings is limited by the number of donors we analyzed in this study, our data reveal a number of novel findings.

Iso–RBD antibodies have been sparsely observed in peripheral B cell analysis (91, 92, 113) and have not been documented in serological analyses. For the first time, we demonstrate the high prevalence of iso–RBD antibodies in serum following infection compared to the infection-naïve, vaccinated group (**Fig. 1**, **D** and **E**). While it is worth noting that the vaccinated group consisted of individuals with Cystic Fibrosis, studies have shown that individuals with Cystic Fibrosis respond similarly to COVID-19 vaccines as healthy individuals (114–116). Our observation of the infection-naïve, vaccinated individuals lacking iso–RBD antibodies suggests that these antibodies are likely elicited by the dissociated S1 during the membrane fusion process in infection (3, 4, 111). Furthermore, our data showing that the iso–RBD mAb, 3548-RBD-15, is able to bind to the RBD of detached S1 further supports this notion (**Supplementary Table 6**) and validates the proteomic identification of iso–RBD antibodies in serum.

While 3548-RBD-15 showed robust ADCP, ADCC, and ADCD activities when tested in the context of immune complexes forming with just RBD, it completely lacked any Fc-mediated effector functions, binding, or neutralization activities in the context of prefusion conformation of S (**Fig. 4**). Iso–RBD antibodies were also less likely to be cross-reactive to SARS-CoV-2 B.1.351 (**Fig. 3F**), which contains conserved mutations, including K417N which is more exposed after S1 is cleaved. It is worth noting that recent VoCs, XBB.1.5 and BQ.1.1, also contain this mutation (117). Based on these results, it is unlikely iso–RBD antibodies can provide protection *in vivo* against infection. In fact, this shedding of S1 during the fusion process may help SARS-CoV-2 misdirect antibody responses, resulting in the elicitation of non-protective antibodies and immune evasion of the virus. This is particularly important in light of ongoing efforts on using RBD as immunogens (10–15) which may elicit potentially non-protective iso–RBD antibodies. We also observed the increased elicitation of S–RBD antibodies in the individuals vaccinated with prefusion-stabilized S as immunogens (**Fig. 1F**). This is important as S–RBD antibodies are likely to be neutralizing (16–18); among our panel of mAbs, the two S–RBD mAbs, 3546-S-2 and 3548-S-2, showed the highest neutralization activity (**Fig. 4B**). Taken together, the use of prefusion-stabilized S as a vaccine immunogen might have had the unintended but beneficial impact of minimizing the elicitation of iso–RBD antibodies while improving elicitation of S–RBD antibodies.

Endogenous hCoVs are relatively benign respiratory pathogens and the second most frequent cause of the common cold (118). Due to their ubiquitous nature, large proportions of the population are exposed to these hCoVs early in childhood (119, 120), and it has been shown that a high fraction of adults have serum IgG titers against endogenous hCoVs (121). In subject-3548, the most abundant serum IgG clonotype, clonotype-27239, targeted a conserved epitope recognizing S from variants (B.1.351 and B.1.1.529) as well as HKU1 (**Fig. 4A**). Combined with its high level of SHM observed at M1 and its high abundance following infection (**Fig. 3H**), this B cell lineage was likely established before the pandemic. These features were also similar to serum IgG clonotypes that comprised the serological memory and persisted in circulation in the context of influenza (80). These findings highlight the potential impact one’s exposure history can have on antibody responses to subsequent vaccination or infection. While the representative mAb from clonotype-27239, 3548-S-1, lacked neutralization activity, it induced robust Fc-mediated effector functions to SARS-CoV-2 as well as HKU1; it remains to be determined whether it can provide robust protection *in vivo*. As the battle with viral evolution continues to persist, understanding the impact of memory-derived antibody repertoire generated by prior hCoVs exposures in mounting effective immune responses to SARS-CoV-2 VoCs will be critical.

We also observed that the composition of the serological IgG repertoire to S stayed constant over 6 months following infection (**Fig. 2 H-J**), although the total titers rapidly decayed for each of the four donors analyzed in our study (**Fig. 2**, **A** and **B**). It has been shown that B cells continue to mature many months following SARS-CoV-2 infection (20, 89, 90), and others have observed that, particularly in individuals with severe COVID-19, germinal center reactions continue to drive antibody evolution (122). Correspondingly, serum antibodies from subject-3548 are more similar to those seen in influenza-vaccinated adults (**Fig. 1B**) than from the other subjects. As serum neutralizing titers decreased more substantially than binding titers (**Fig. 2 A-C**) and non-IgG isotypes contributed mainly to the M1 and M3 neutralizing titer (**Fig. 2 E**), a distinct but shorter-lived IgM and IgA response likely is essential to the early convalescent protective antibody response. Indeed, many others have noted that IgA is critical for protection (106), although the titer wanes rapidly following infection (27, 106).

As we demonstrate in our work, the elicitation of iso–RBD antibodies may be critical to consider when designing new immunogens, as they are likely to not recognize S on the viral surface. Over 70 million individuals contracted COVID-19 before the release of the first FDA-approved vaccine, resulting in a large population being vaccinated after infection. It will be critical to understand the vaccine response in previously infected individuals and to track the contribution of antibodies originally elicited from infection compared to those elicited from the following vaccinations. Additionally, elucidating serum antibodies that originate from pre-pandemic will be necessary to understand the contribution of hCoV-elicited serum antibodies in the response to SARS-CoV-2 infection and vaccinations as we have observed these antibodies may contribute a significant fraction to the serological response.

## Materials and Methods

### Sample Collection

All study protocols were approved by the Institutional Review Boards at Dartmouth College and Dartmouth-Hitchcock Medical Center. Informed consent was obtained from the participants. Blood was taken after (M1 between 31 and 46 days, M3 between 91 and 97 days, and M6 between 184 and 201 days) infection for the SARS-CoV-2 infected donors. Infected donor samples were collected under study number 020082. In our vaccinated donors with cystic fibrosis, blood was taken after (both 7 and 28 days) the second SARS-CoV-2 vaccination (either with BNT162b2 or mRNA-1273 vaccination) in a two-dose series. Vaccinated donor samples were collected under study number 02000827.

### High-throughput Sequencing of V_H_ and V_L_

500 ng of total RNA from peripheral blood mononuclear cells (PBMCs) were taken 1-month post-infection and reverse transcribed according to the manufacturer’s instructions using SuperScript IV enzyme (Invitrogen) and Oligo(dT) primer (Invitrogen). V_H_ and V_L_-only transcripts were amplified using the FastStart High Fidelity PCR System (Roche) with gene-specific primers (123). V_H_ and V_L_ amplicons were sequenced using the Illumina MiSeq platform. Sequences were annotated using MiXCR 2.1.5. IMGT library v3 was used to annotate V_H_ and V_L_ sequences.

### V_H_:V_L_-paired Sequencing

Paired heavy and light chain sequencing of B cells from each donor’s M1 post-infection B cells was conducted as done previously (79, 80). Briefly, PBMCs were isolated as single cells inside emulsion droplets using a custom flow-focusing apparatus. Droplets contained lysis buffer and poly(dT) conjugated magnetic beads to capture heavy and light chain mRNA. Magnetic beads were collected and emulsified to serve as a template for emulsion overlap extension RT-PCR, in which V_H_ and V_L_ amplicons were physically linked. The V_H_:V_L_ amplicons were extracted from the emulsions, amplified with a nested PCR, and sequenced using the Illumina MiSeq platform. V_H_ and V_L_ regions were amplified separately for full-length V_H_ and V_L_ analysis for the recombinant expression of monoclonal antibodies using the Illumina MiSeq platform as previously described (79, 80).

### Preparation of Total IgG and IgA from Sera

Each serum sample analyzed in this study was first passed through a 1 mL Protein G agarose (Thermo Fisher, 20397) affinity column in gravity mode. Serum flow-through was collected and passed through the column three times. The column was washed with 20 mL of PBS prior to elution with 5 mL of 100 mM glycine-HCl, pH 2.7. The eluate, containing total IgG from serum, was immediately neutralized with 0.75 mL of 1 M Tris-HCl, pH 8.0. Serum flow-through was then passed through 1 mL of Peptide M agarose (InvivoGen, gel-pdm-5) affinity column in gravity mode. Serum flow-through was collected and passed through the column three times. The column was washed with 20 mL of PBS prior to elution with 5 mL of 100 mM glycine-HCl, pH 2.7. The eluate, containing total IgA from serum, was immediately neutralized with 0.75 mL of 1 M Tris-HCl, pH 8.0. Purified IgG was digested into F(ab’)2 with 25 μg of recombinantly-made IdeS protease per 1 mg of IgG for 4 hr on a rotator at 37°C and then incubated with Strep-Tactin agarose (IBA-Lifesciences, 2-1206-025) for 1 hr to remove IdeS protease.

### Antigen-enrichment and MS Sample Preparation

SARS-CoV-2 S (Wu, GenBank_MN908947, residues 14 to 1208 with HexaPro (R682G, R683S, A684A, R685S, K986P, V987P, F817P, A892P, A899P, and A942P mutations)) (84) and hCoV-HKU1 S (GenBank_NC006577, residues 13 to 1276 with R752G, R753G, K754S, R755G, R756R, N1067P, and L1068P mutations) were recombinantly expressed on Expi293F cells (Invitrogen) and purified by HisPur™ Cobalt resin (Thermo Fisher, 89966). SARS-CoV-2 RBD (Wu, S residues 319 to 591) and SARS-CoV-2 RBD (B.1.351, S residues 319 to 591 with K417N, E484K, and N501Y mutations from Wu strain) were recombinantly expressed on Expi293F cells and purified by Protein G agarose. Recombinant antigen was immobilized on N-hydroxysuccinimide (NHS)–activated agarose resins (Thermo Fisher, 26197) by overnight rotation at 4°C. The coupled agarose resins were washed with PBS, and unreacted NHS groups were blocked with 1 M Tris-HCl, pH 7.5 for 30 min at RT. The resins were further washed with PBS and packed into a 0.8 mL centrifuge column (Thermo Fisher, 89868).

For each sample, F(ab’)2 was incubated with the individual antigen affinity columns for 1 hr on a rotator at RT. Flow-through was collected, and the column was washed with 5 mL of PBS. Antigen-enriched F(ab’)2 was eluted with 1% (v/v) formic acid in 0.5 mL fractions. 30 μL from each elution fraction, neutralized with 1M Tris-HCl, pH 8.0, and pre-flow-through, flow-through, and wash samples were assayed by indirect ELISA with each antigen. ELISA signal in each elution sample was checked using 1:5000 diluted goat anti-human IgG (Fab specific) HRP-conjugated secondary antibodies (Sigma-Aldrich, A0293-1ML). Elution fractions showing an ELISA signal were pooled and concentrated under vacuum to a volume of ∼1 μL. 40 μL of DPBS and 2 μL of Tris-HCl, pH 7.5 were added before neutralizing with 1 M NaOH. The neutralized elution samples were concentrated to 50 μL under vacuum.

For each enrichment, elution and flow-through samples were denatured with 50 μL of 2,2,2-trifluoroethanol (TFE) and 5 μL of 100 mM dithiothreitol (DTT) at 55°C for 1 hr, and then alkylated by incubation with 3 μL of 550 mM iodoacetamide (Sigma) for 30 min at RT in the dark. Alkylation was quenched with 892 μL of 40 mM Tris-HCl, and protein was digested with trypsin (1:30 (w/w) trypsin/protein) for 16 hrs at 37°C. Formic acid was added to 1% (v/v) to quench the digestion, and the sample volume was reduced to ∼150 μL under vacuum. Peptides were then bound to a Hypersep SpinTip C-18 (Thermo Scientific, 60109-412), washed three times with 0.1% formic acid, and eluted with a 60% acetonitrile and 0.1% formic acid solution. C18 eluate was concentrated under vacuum centrifugation and resuspended in 50 μL in 5% acetonitrile, 0.1% formic acid.

### LC-MS/MS Analysis

Samples were analyzed by liquid chromatography-tandem mass spectrometry on an Easy-nLC 1200 (Thermo Fisher Scientific) coupled to an Orbitrap Fusion Tribrid (Thermo Scientific). Peptides were first loaded onto an Acclaim PepMap RSLC NanoTrap column (Dionex; Thermo Scientific) prior to separation on a 75 μm × 15 cm Acclaim PepMap RSLC C18 column (Dionex; Thermo Scientific) using a 1.6%–76% (v/v) acetonitrile gradient over 90 mins at 300 nL/min. Eluting peptides were injected directly into the mass spectrometer using an EASY-Spray source (Thermo Scientific). The instrument was operated in data-dependent mode with parent ion scans (MS1) collected at 120,000 resolution. Monoisotopic precursor selection and charge state screening were enabled. Ions with charge ≥ +2 were selected for collision-induced dissociation fragmentation spectrum acquisition (MS2) in the ion trap, with a maximum of 20 MS2 scans per MS1. Dynamic exclusion was active with a 15-s exclusion time for ions selected more than twice in a 30-s window. Each sample was run three times to generate technical replicate datasets.

### MS/MS Data Analysis

Protein sequence databases were constructed using the V_H_ and V_L_ sequences obtained from each donor. V_H_ and V_L_ sequences with ≥ 2 reads were concatenated to a database of background proteins comprising a consensus human protein database (Ensembl 73, longest sequence/gene) and a list of common protein contaminants (MaxQuant). Spectra were searched against the database using SEQUEST (Proteome Discoverer 2.4; Thermo Scientific). Searches considered fully tryptic peptides only, allowing up to two missed cleavages. A precursor mass tolerance of 5 ppm and fragment mass tolerance of 0.5 Da were used. Modifications of carbamidomethyl cysteine (static) and oxidized methionine (dynamic) were selected. High-confidence peptide-spectrum matches (PSMs) were filtered at a false discovery rate of <1% as calculated by Percolator (q-value <0.01, Proteome Discoverer 2.4; Thermo Scientific).

Iso/Leu sequence variants were collapsed into single peptide groups. For each scan, PSMs were ranked first by posterior error probability (PEP), then q-value, and finally XCorr. Only unambiguous top-ranked PSMs were kept; scans with multiple top-ranked PSMs (equivalent PEP, q-value, and XCorr) were designated ambiguous identifications and removed. The average mass deviation (AMD) for each peptide was calculated as described (77) using data from elution only. Peptides with AMD > 1.7 ppm were removed. Additionally, only peptides identified in ≥ 2 injections were kept as high-confidence identifications.

Peptide abundance was calculated from the extracted-ion chromatogram (XIC) peak area, as described (78). For each peptide, a total XIC area was calculated as the sum of all unique peptide XIC areas of associated precursor ions. The average XIC area across replicate injections was calculated for each sample. For each antigen dataset, the eluate and flow-through abundances were compared and peptides with ≥5-fold higher signal in the elution sample were considered to be antigen-specific.

### Peptide-to-Clonotype Index and Mapping

V_H_ sequences were grouped into clonotypes based on hierarchical clustering as previously described (78). Cluster membership required ≥90% identity across the CDRH3 amino acid sequence as measured by the edit distance. High-confidence peptides identified by MS/MS analysis were mapped to clonotypes, and peptides uniquely mapping to a single clonotype were considered ‘informative’. The abundance of each antibody clonotype was calculated by summing the XIC areas of the informative peptides mapping to ≥5 amino acids of the CDRH3 region. SHM levels for individual clonotypes were calculated by averaging the SHM rates of all the V_H_ sequences within each clonotype that contained the detected CDRH3 sequences.

### Recombinant mAb Expression and Purification

Selection of antibody sequences for recombinant expression was based on the combination of V_H_:V_L_-paired databases and proteomics data. First, we identified antibody clonotypes identified in the proteomics analysis and searched for the same clonotype in the V_H_:V_L_-paired database. Full-length heavy and light chain sequences were then determined from the paired sequencing database. These genes were purchased as eBlocks (Integrated DNA Technologies) and cloned into the pcDNA3.4 vector (Invitrogen). Heavy and light chain plasmids for each monoclonal antibody were transfected into Expi293 cells at a 1:3 ratio. After incubating for 5 days at 37°C with 8% CO2, the supernatant containing secreted antibodies was collected by centrifugation at 4000 rpm for 15 min at 4°C. Supernatant was passed over a column with 1.0 mL Protein G agarose resin three times to ensure efficient capture. After washing the column with 20 mL of PBS, antibodies were eluted with 5 mL 100 mM glycine-HCl, pH 2.7 and immediately neutralized with 0.75 mL 1 M Tris-HCl, pH 8.0. Antibodies were buffer exchanged into PBS utilizing Amicon Ultra-30 centrifugal spin columns (Millipore).

## ELISAs

50% effective concentration (EC_50_) values and AUC values based on ELISA were used to determine the binding apparent affinities of the recombinant monoclonal antibodies and serum. SARS-CoV-2 S (Wu, residues 14 to 1213) was purchased from SinoBiological (40589-V08B1) and Biolegend (793704). The following reagents were obtained through BEI Resources, NIAID, NIH: Spike Glycoprotein (Stabilized) from SARS-Related Coronavirus 2, Wuhan-Hu-1 with C-Terminal Histidine and Twin-Strep^®^ Tags, Recombinant from HEK293 Cells, NR-52724, Spike Glycoprotein (Stabilized) from SARS-Related Coronavirus 2, Wuhan-Hu-1 with C-Terminal Histidine and Twin-Strep^®^ Tags, Recombinant from CHO Cells, NR-53937, and SARS-CoV-2 Spike S1-His Tag (HPLC-verified) (HEK293 cells expressed) SinoBio Cat: 40591-V08H (fine purification), NR-53798. SARS-CoV-2 S B.1.1.529 was purchased from SinoBiological (40589-V08H26). SARS-CoV-2 S, SARS-CoV-2 RBD, SARS-CoV-2 B.1.351. RBD, SARS-CoV-2 B.1.351. S (Residues 14 to 1208, L18F, D80A, D215G, Δ242-244, R246I, K417N, E484K, N501Y/T, D614G, A701V mutations from Wu-Hu-1 strain) and hCoV-HKU1 S were recombinantly expressed on Expi293F (Invitrogen) and purified by HisPur™ Cobalt resin (Thermo Fisher, 89966). First, costar 96-well ELISA plates (Corning) were coated overnight at 4°C with 4 μg/mL recombinant antigens, washed with PBST (0.01% Tween20 in PBS), and blocked with 0.5% BSA in PBS for 1 hr at RT. After blocking, serially diluted recombinant antibodies or serum samples bound to the plates for 1 hr, followed by 1:5000 diluted goat anti-human IgG (Fab-specific) HRP-conjugated secondary antibodies (Sigma-Aldrich, A0293-1ML) for 1 hr. For detection, 100 μL TMB-ELISA substrate solution (Thermo Scientific, 34028) was added before quenching with 50 μL 1 M H_2_SO_4_. Absorbance was measured at 450 nm using a SpectraMax i3x plate reader. Data were analyzed and fitted for EC_50_ using a 4-parameter logistic nonlinear regression model in the GraphPad Prism 10 software.

### Biolayer Interferometry (BLI)

K_D_, *k*_on,_ and *k*_off_ of recombinant mAbs were measured by BLI using a ForteBio Octet RED96 (Molecular Devices). Antigens and antibodies were diluted in PBSF (PBS with 0.1% w/v BSA), and assays were run at 25°C at 1000 rpm orbital shaking speed. Streptavidin tips (SAX2, Molecular Devices) were first incubated in PBSF for thirty minutes before being bound to biotinylated 6x-His Tag Monoclonal antibodies (Thermo Scientific, MA1-21315-BTIN) until a binding threshold of 0.2 nm was met for any tip. For each binding assay, after a 360 s baseline step in PBSF, tips were dipped into 100 mM of his-tagged antigen for 360 s and then dipped into PBSF for 360 s to measure the dissociation of antigen from the biosensor. Tips were then dipped in antibody solution (ranging from 500 nM to 2.6 nM) for 360 s, and dissociation was measured in PBSF for 360 s. Data was aligned to the start of the association step, corrected using Savitzky-Golay filtering and fit to an association/dissociation model using GraphPad PRISM.

### Pseudovirus Neutralization Assays

Samples of total serum and serum IgG from SARS-CoV-2 infected donors and recombinant mAbs were tested in microneutralization assays using a VSV-SARS-CoV pseudovirus system (83, 124). Serum IgG was purified using the Melon Gel IgG Spin Purification kit (Thermo Fisher, 45206) following the manufacturer’s instructions. Samples were serially diluted 2-fold (1:12.5-1:6400 for total serum; 1:25-1:1600 for serum IgG; 37.5 μg/mL to 0.1 μg/mL for recombinant mAbs; 1.0 μg/mL to 7.8 ng/mL for control mAbs) and incubated with VSV-SARS-CoV-2 S or VSV-SARS-CoV-2 B.1.351 S pseudovirus for 1 hr at 37°C before adding to 293T-hsACE2-expressing cells (Integral Molecular). Plates were incubated at 37°C, 5% CO_2_ for 24 hr, after which luciferase activity was measured in cell lysates using the Bright-Glo system (Promega) with a Bio-Tek II plate reader. Percent neutralization was calculated as 100 - (mean RLU test wells/mean RLU positive control wells) x 100 and used to determine the 50% neutralization titers for serum (NT50) and half-maximal inhibitory concentrations for monoclonal antibodies (IC50).

### Fc-mediated Functional Assays

Antibody-dependent cellular phagocytosis (ADCP) by monocytes was performed essentially as described (125, 126). Briefly, 500,000 1 μm yellow-green fluorescent microspheres (Thermo, F8813) were covalently conjugated with recombinant SARS-CoV-2 S, SARS-CoV-2 RBD, SARS-CoV-2 S (B.1.351), SARS-CoV-2 RBD (B.1.351), and hCoV HKU1 S and incubated for 3 hr with diluted recombinant mAbs (3-fold dilution, 5 μg/mL to 6.8 ng/mL) and 20,000 cells from the human monocytic THP-1 cell line (ATCC, TIB-202). After pelleting, washing, and fixing, phagocytic scores were quantified as the product of the percentage of cells that phagocytosed one or more fluorescent beads and the median fluorescent intensity of this population as measured by flow cytometry with a MACSQuant Analyzer (Miltenyi Biotec). ADCP assays were performed in triplicate with high correspondence between results presented here and the replicate run. REGN10987 IgG1 and REGN10987 IgG2σ (127) were used as positive and negative controls, respectively. Wells containing no antibody were used to define the level of antibody-independent phagocytosis.

Antibody-dependent cellular cytotoxicity (ADCC) potential was measured using a Jurkat Lucia NFAT cell line (Invivogen, jktl-nfat-cd16), cultured according to the manufacturer’s instructions, in which engagement of FcγRIIIa (CD16) on the cell surface leads to the secretion of luciferase (82, 83). The high binding 96 well plate was coated with 50uL of 1 μg/mL SARS-CoV-2 S, SARS-CoV-2 RBD, SARS-CoV-2 S (B.1.351), SARS-CoV-2 RBD (B.1.351), and hCoV HKU1 S overnight at 4°C. Plates were then washed with PBST and blocked with 2.5% BSA in PBS for 1 hr at RT. After washing, diluted recombinant mAbs (3-fold dilution, 5 μg/mL to 6.8 ng/mL) and 1 × 10^5^ cells per well in growth medium lacking antibiotics were cultured at 37°C for 24 hr in a 200 μL volume. A 25 μL volume of supernatant was drawn from each well and transferred to an opaque, white 96 well plate, to which 75 μL of QuantiLuc substrate was added and luminescence immediately read on a SpectraMax Paradigm plate reader (Molecular Devices) using 1 s of integration time. The reported values are the mean of three kinetic reads taken at 0-, 2.5-, and 5-min. Negative control wells substituted assay medium for sample while 1x cell stimulation cocktail (Thermo, 00-4970-93) plus an additional 2 mg/mL ionomycin were used to induce expression of the transgene as a positive control.

Antibody-dependent complement deposition (ADCD) was quantified essentially as previously described (128). In brief, diluted recombinant mAbs (3-fold dilution, 5 μg/mL to 6.8 ng/mL) were incubated at RT for 2 hr with multiplex assay microspheres. After washing, each sample was incubated with human complement serum (Sigma, S1764) at a concentration of 1:50 at RT with shaking for 1 hr. Samples were washed, sonicated, and incubated with murine anti-C3b (Cedarlane, CL7636AP) at RT for 1 hr followed by anti-mouse IgG1-PE secondary antibody (Southern Biotech, 1070-09) at RT for 30 min. After a final wash and sonication, samples were resuspended in Luminex sheath fluid, and complement deposition was determined on a FlexMax 3D Array Reader (Luminex Corp) instrument to define the MFI. Assays performed with no antibody and with heat-inactivated human complement serum were used as negative controls.

### Statistical Analysis

All statistical analyses were performed using GraphPad Prism 10.0 (GraphPad Software, San Diego, CA). All the statistical tests performed are described in the figure legends, and correlations were considered significant at a *p*-value of < 0.05 (\**p* < 0.05, \*\**p* < 0.01, \*\*\**p* < 0.001, n.s.; not significant).

### Data Availability

The raw proteomic data and high-throughput sequences have been deposited in MassIVE (https://massive.ucsd.edu/ProteoSAFe/static/massive.jsp) under accession ID MSVXXXXXXXXX. Nucleotide sequences of individual monoclonal antibodies characterized in this paper are deposited at GenBank (accession numbers GenBank: OQXXXXXX-OQXXXXXX).

## Acknowledgements

The research was supported by NIH grant number P20 GM113132 (to J.L.), the Cystic Fibrosis Foundation grant number STANTO19R0 (to J.L.), NIH grant number R01 AI146779 (to A.G.S.), a Massachusetts Consortium on Pathogenesis Readiness (MassCPR) grant (to A.G.S.), NSF grant number 1840344 (to N.C.C.), NIGMS T32 GM007753 (to B.M.H.), T32 AI007245 (to J.F.), and F30 AI160908 (to B.M.H.). The authors also acknowledge the support from bioMT at Dartmouth through NIH NIGMS Grant P20 GM113132 and Immune Monitoring and Flow Cytometry Resource (IMFCSR) at the Norris Cotton Cancer Center at Dartmouth through NCI Cancer Center Support Grant 5P30 CA023108-41.

---

**Supplementary Figure 1.**
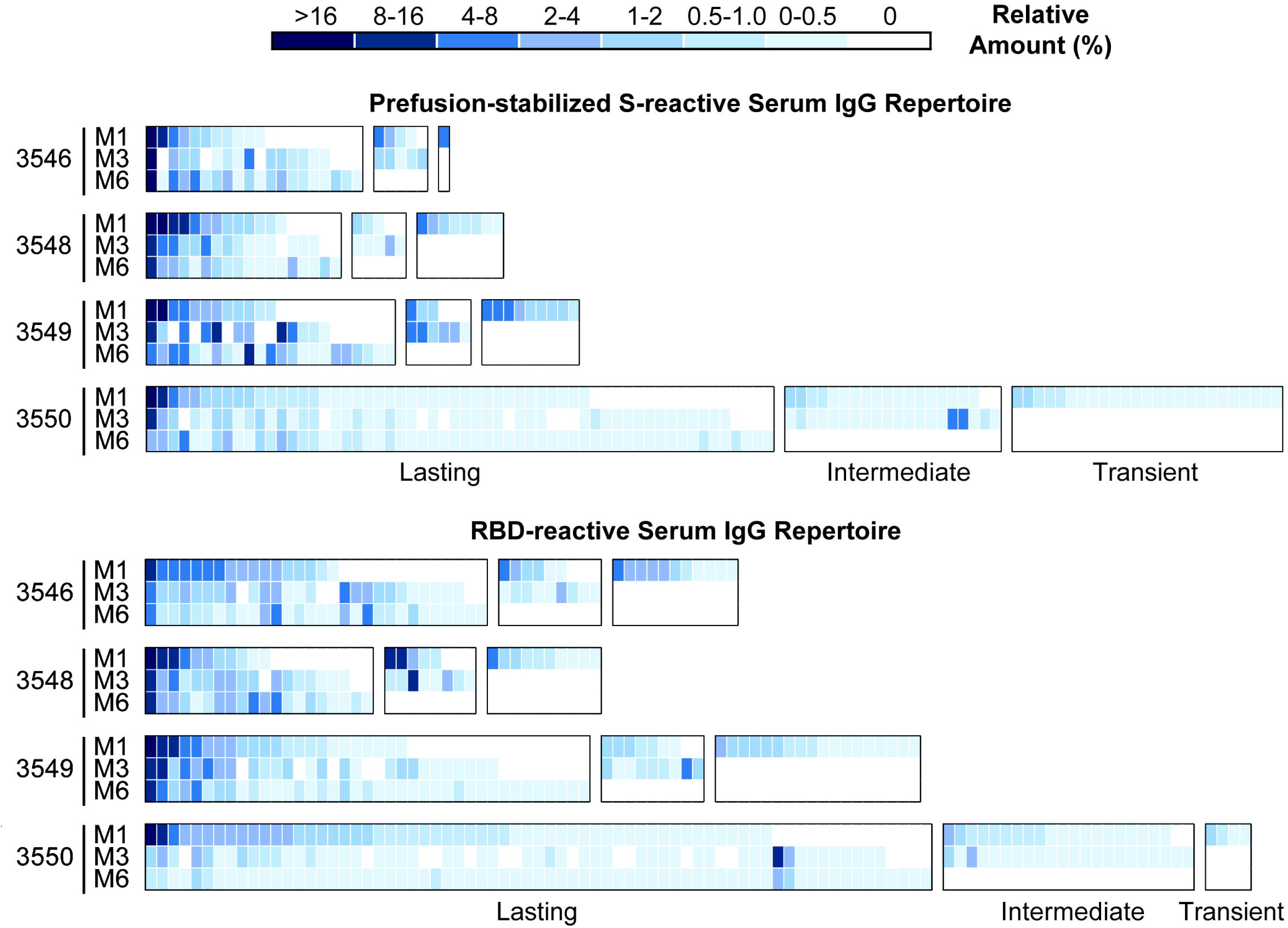
Serological repertoire of subject-3546, subject-3548, subject-3549, and subject-3550. Each column is a unique clonotype. The relative amount of each clonotype is calculated by multiplying its relative abundance by the serum binding titer for that sample before normalizing it to the M1 titer.

**Supplementary Figure 2.**
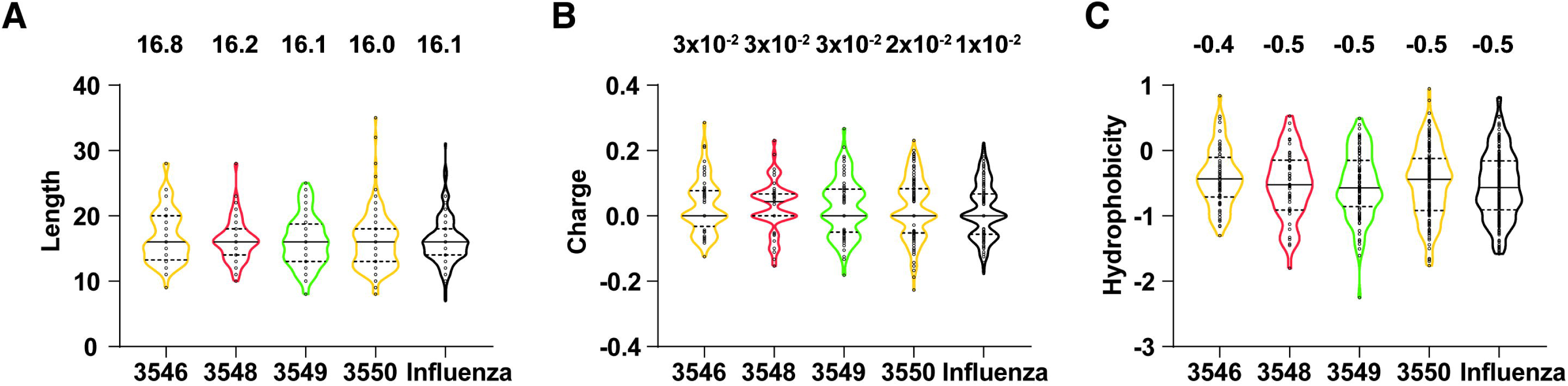
Comparison of CDRH3 length. (**A**), charge (**B**), and hydrophobicity (**C**) of the anti-SARS-CoV-2 S serological repertoire. The influenza data set is based on previously published HA-reactive serum antibodies identified in four adults following trivalent influenza vaccination (79). Hydrophobicity values are based on the Kyte-Doolittle scale. Black horizontal lines represent medians (solid) and quartiles (dotted), with mean values included above each plot.

**Supplementary Figure 3.**
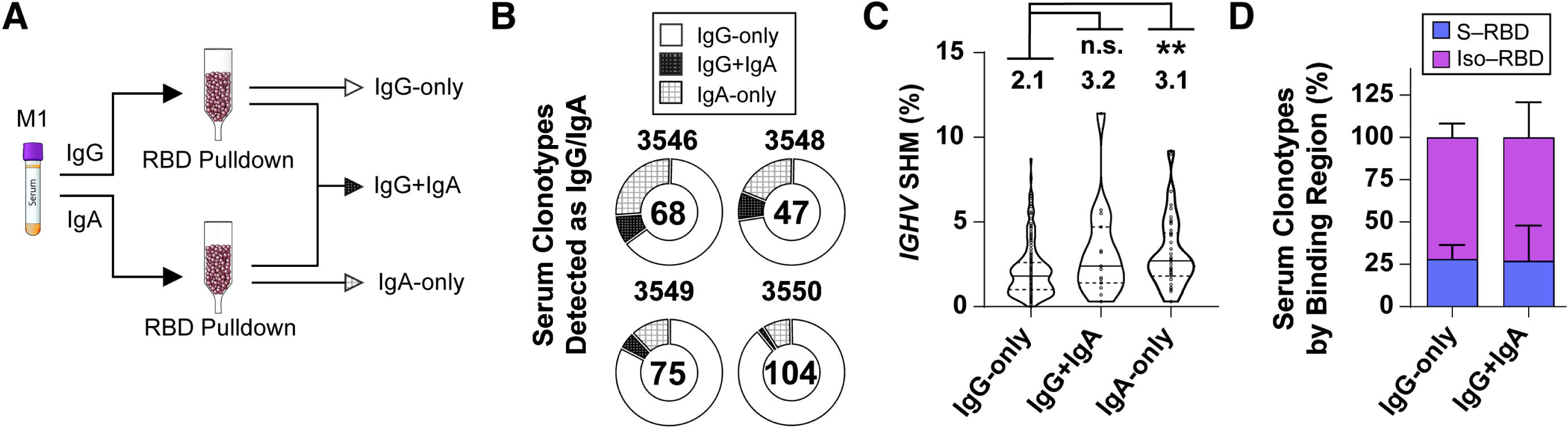
Profiling the serum IgA repertoire to RBD. **A.** Experimental design. IgG and IgA specific to RBD in M1 serum samples was isolated through RBD-immobilized affinity purification, and purified IgG and IgA were subsequently analyzed proteomically. **B.** Clonotypic diversity of serum antibody clonotypes that are either only detected as IgG (IgG-only) or IgA (IgA-only) and those detected as both IgG and IgA (IgG+IgA). The total number of clonotypes is included at the center of each plot. **C.** *IGHV* SHM level of IgG-only, IgA-only, IgG+IgA clonotypes. Black horizontal lines represent medians (solid) and quartiles (dotted), with mean values included above each plot. Statistical significance is determined by Dunn’s multiple comparisons test. (** *p* < 0.01). **D.** Breakdown of IgG-only and IgG+IgA clonotypes by their binding regions. The error bars indicate SD.

**Supplementary Figure 4.**
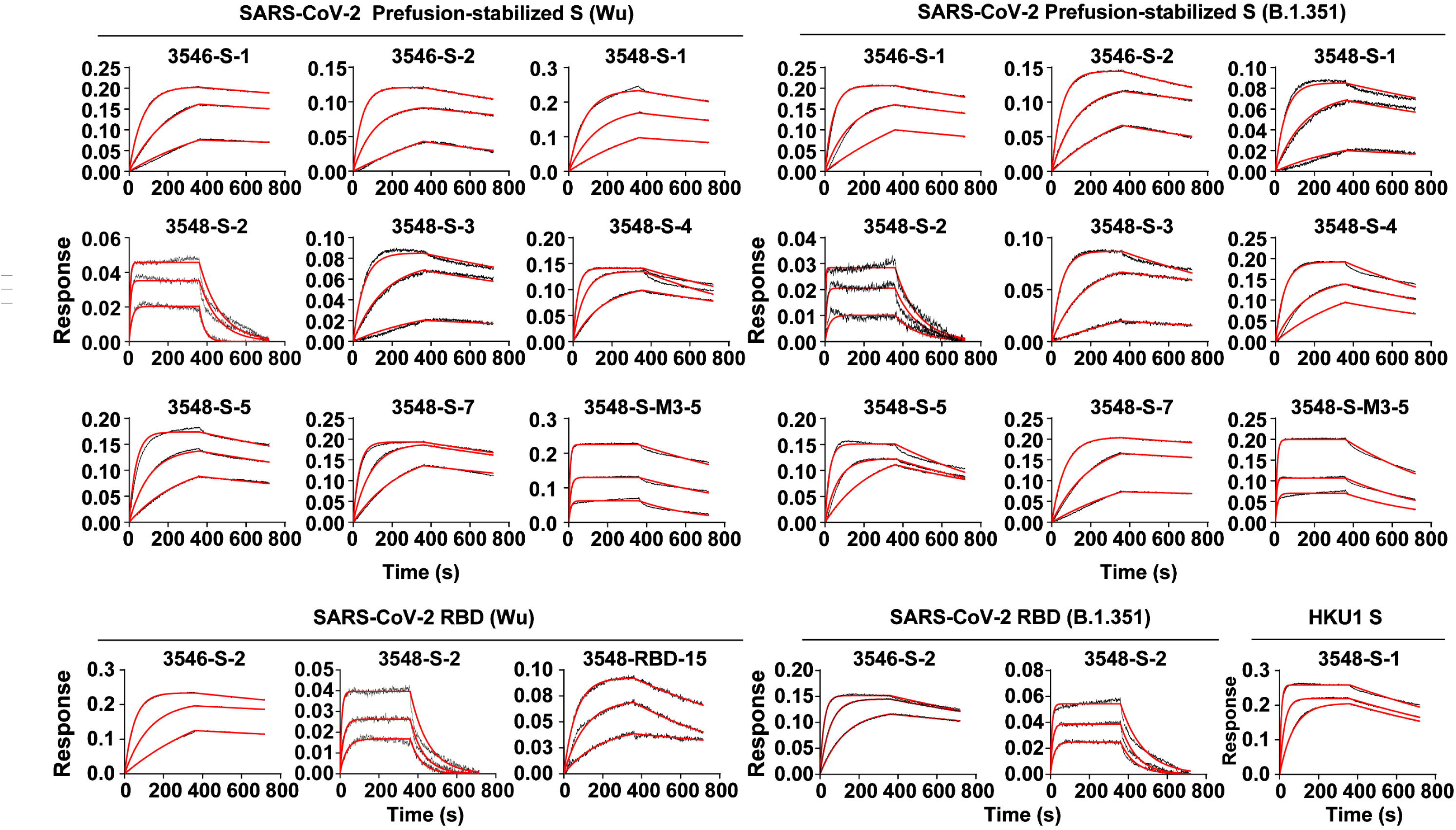
Biolayer interferometry binding data of selected mAbs. Assays are run at three concentrations (18.8 nM, 6.3 nM, and 2.1 nM for all mAbs except for 3548-S-2 and 3548-RBD-15 which were run at 500 nM, 166.6 nM, and 55.5 nM), and binding kinetics were determined using GraphPad PRISM constraining each concentration to have independent *k*_on_ and *k*_off_ values.

**Supplementary Figure 5.**
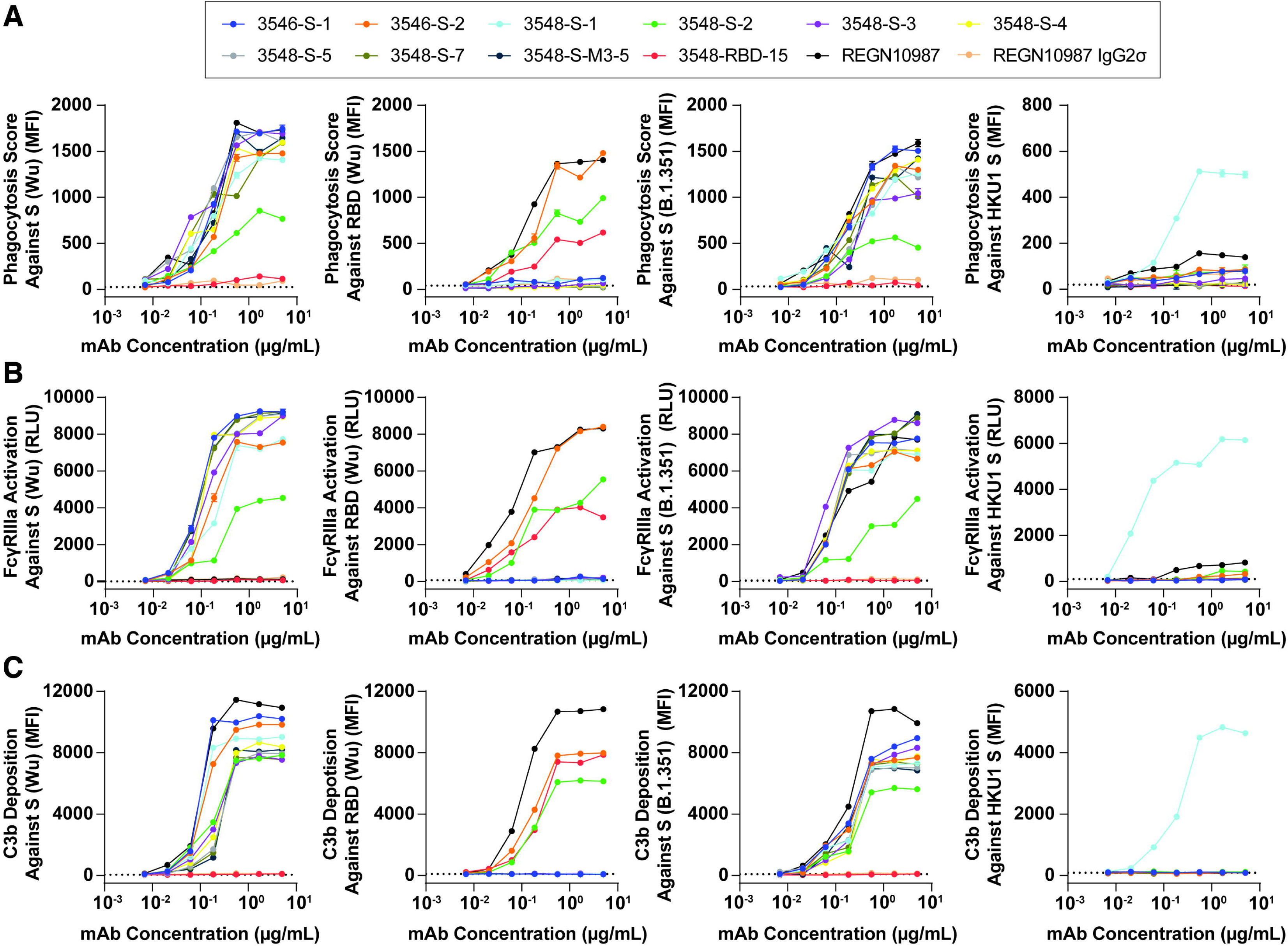
Fc-mediated functional activity of SARS-CoV-2 mAbs. Antibody-dependent cellular phagocytosis (**A**), FcγRIIIa activation (**B**), and antibody-dependent complement deposition (**C**) activity of mAbs against SARS-CoV-2 prefusion-stabilized S (Wu), RBD (Wu), prefusion-stabilized S (B.1351), and HKU1 S. The dotted line indicates the no antibody control, and REGN10987 IgG2σ (127) was used as a null Fc-mediated effector function control.

**Supplementary Figure 6.**
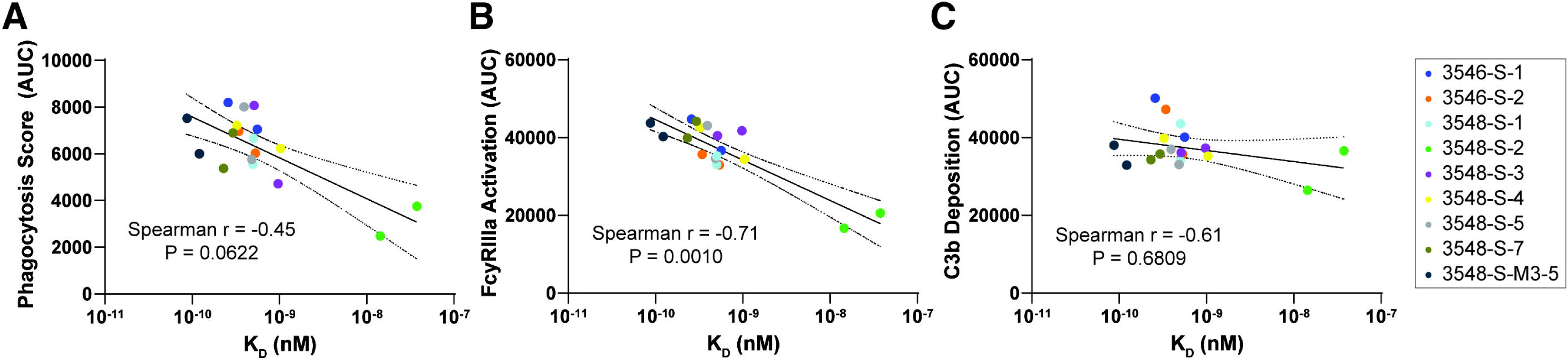
Correlation between functional activity and binding kinetics of SARS-CoV-2 mAbs, Antibody-dependent cellular phagocytosis (**A**), FcγRIIIa activation (**B**), and antibody-dependent complement deposition (**C**) activity of mAbs targeting SARS-CoV-2 prefusion-stabilized S (Wu), and prefusion-stabilized S (B.1351) were plotted with their corresponding K_D_ values from BLI. Correlation was tested using a Spearman correlation test. Spearman *r* and *P* values are displayed on each plot. Semi-log lines are drawn as solid lines with 95% confidence intervals represented by dotted lines.

**Supplementary Table 1.** Subject and sample summary.

| Subject | Age | Severity | Date of Illness / Positive Test | Date of M1 Blood Draw | Date of M3 Blood Draw | Date of M6 Blood Draw |
| --- | --- | --- | --- | --- | --- | --- |
| 3546 | 62 | Moderate | 03/07/20 | 04/13/20 | 06/08/20 | 9/24/20 |
| 3548 | 63 | Severe | 03/15/20 | 04/15/20 | 06/15/20 | 9/15/20 |
| 3549 | 59 | Mild | 03/11/20 | 04/15/20 | 06/16/20 | 9/16/20 |
| 3550 | 52 | Moderate | 03/08/20 | 04/23/20 | 06/17/20 | 9/16/20 |

**Supplementary Table 2.** Summary of Next-Gen Sequencing of peripheral B cells isolated at M1.

| Subject | B Cell Count | Total Number of Sequence Reads | Number of Unique Antibody Sequences |
| --- | --- | --- | --- |
| 3546 | $1.53 \times 10^6$ | 5,943,497 | 477,944 |
| 3548 | $0.78 \times 10^6$ | 6,760,982 | 350,835 |
| 3549 | $1.31 \times 10^6$ | 7,389,287 | 178,876 |
| 3550 | $2.33 \times 10^6$ | 6,303,524 | 188,188 |

**Supplementary Table 3.** Polarization of the serological repertoires. The D60 diversity index refers to the number of the most abundant antibody clonotypes which comprise 60% (by abundance) of the serological repertoire. For each time point and pulldown, the diversity index values were divided by the total number of antibody clonotypes identified to determine the frequency of antibody clonotypes constituting the D60 diversity indices.

| Subject | Prefusion-stabilized S Pulldown |  |  | RBD Pulldown |  |  |
| --- | --- | --- | --- | --- | --- | --- |
|  | M1 | M3 | M6 | M1 | M3 | M6 |
| 3546 | 2/16 (12.5%) | 1/19 (5.3%) | 3/20 (15.0%) | 10/34 (29.4%) | 9/34 (26.5%) | 6/30 (20.0%) |
| 3548 | 4/24 (16.7%) | 4/20 (20.0%) | 3/18 (16.7%) | 5/26 (19.2%) | 4/25 (16.0%) | 5/20 (25.0%) |
| 3549 | 5/24 (20.8%) | 6/18 (33.3%) | 5/23 (21.7%) | 5/48 (10.4%) | 5/35 (14.3%) | 4/39 (10.3%) |
| 3550 | 3/84 (3.6%) | 6/66 (9.1%) | 8/58 (13.8%) | 12/79 (15.2%) | 7/74 (9.5%) | 8/69 (11.6%) |

**Supplementary Table 4.**
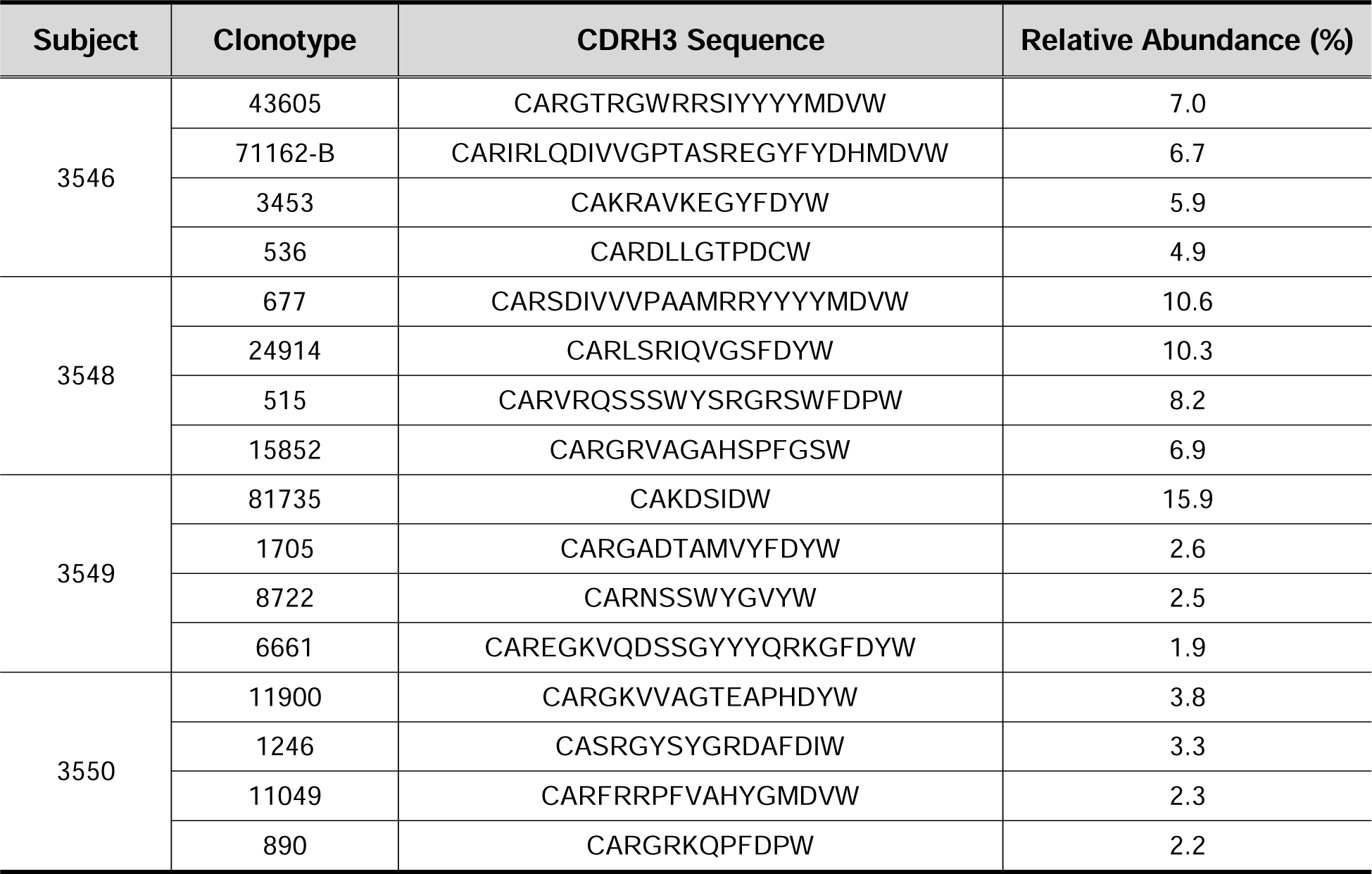
Summary of representative iso–RBD clonotypes. Relative abundance is calculated from the M1 time point.

**Supplementary Table 5.** Summary of the infection-naïve, vaccinated individuals with Cystic Fibrosis.

| Subject | Age | Vaccine Received | Dose 1 Date | Dose 2 Date | Serum Collection Date |
| --- | --- | --- | --- | --- | --- |
| CV-1 | 37 | Pfizer-BioNTech | 3/15/2021 | 4/22/2021 | 5/20/2021 |
| CV-2 | 25 | Pfizer-BioNTech | 3/1/2021 | 3/24/2021 | 4/22/2021 |
| CV-3 | 28 | Moderna | 4/6/2021 | 5/4/2021 | 6/2/2021 |
| CV-4 | 41 | Pfizer-BioNTech | 3/16/2021 | 4/6/2021 | 5/5/2021 |
| CV-5 | 33 | Moderna | 3/29/2021 | 4/26/2021 | 5/24/2021 |
| CV-6 | 19 | Pfizer-BioNTech | 3/13/2021 | 4/2/2021 | 4/30/2021 |
| CV-7 | 60 | Pfizer-BioNTech | 3/18/2021 | 4/15/2021 | 5/13/2021 |
| CV-8 | 31 | Moderna | 3/24/2021 | 4/21/2021 | 5/19/2021 |
| CV-9 | 31 | Pfizer-BioNTech | 4/1/2021 | 4/22/2021 | 5/20/2021 |

**Supplementary Table 6.**
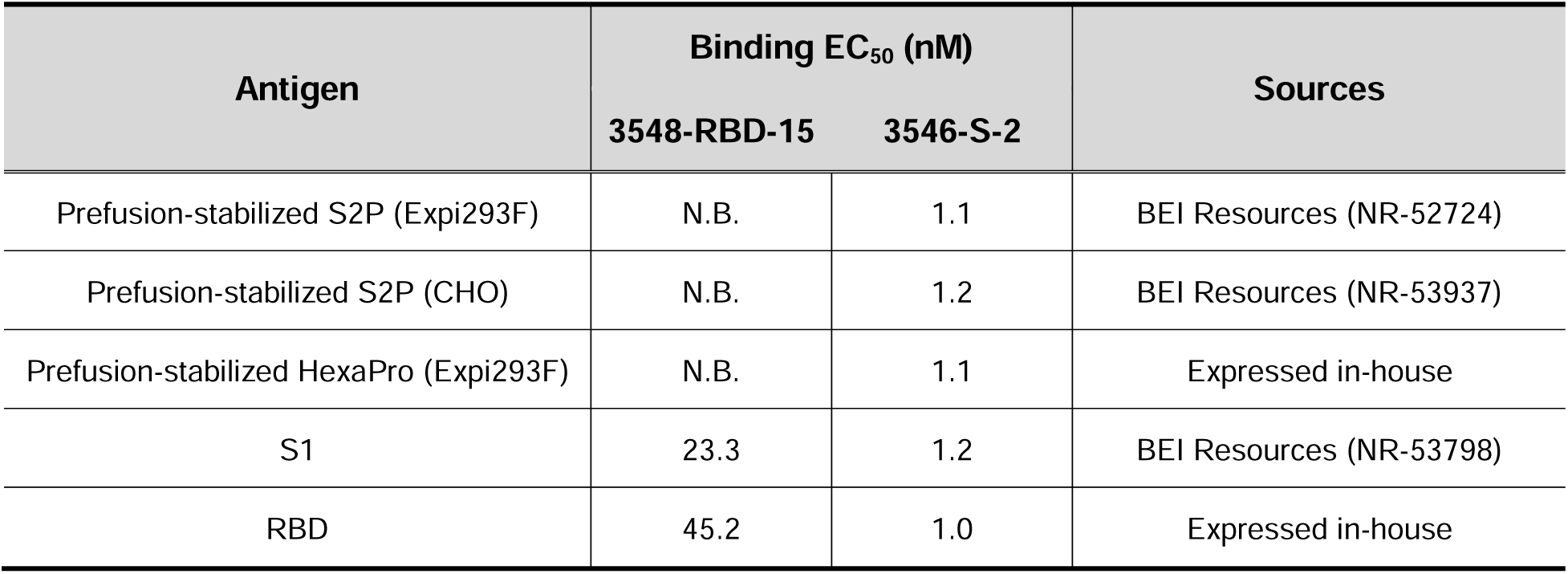
EC_50_ of 3548-RBD-15 and 3546-S-2 against recombinant SARS-CoV-2 S (Wu) proteins from different sources.

| Antigen | Binding EC<sub>50</sub> (nM) |  | Sources |
| --- | --- | --- | --- |
|  | 3548-RBD-15 | 3546-S-2 |  |
| Prefusion-stabilized S2P (Expi293F) | N.B. | 1.1 | BEI Resources (NR-52724) |
| Prefusion-stabilized S2P (CHO) | N.B. | 1.2 | BEI Resources (NR-53937) |
| Prefusion-stabilized HexaPro (Expi293F) | N.B. | 1.1 | Expressed in-house |
| S1 | 23.3 | 1.2 | BEI Resources (NR-53798) |
| RBD | 45.2 | 1.0 | Expressed in-house |

**Supplementary Table 7.** Biolayer interferometry measurements for selected mAbs. The errors represent SD.

| Antigen | mAb ID | $K_D$ (M) | $k_{on}$ ($M^{-1}s^{-1}$) | $k_{off}$ ($s^{-1}$) |
| --- | --- | --- | --- | --- |
| Prefusion-stabilized S (Wu) | 3546-S-1 | $(2.59 \pm 0.02) \times 10^{-10}$ | $(7.76 \pm 0.03) \times 10^5$ | $(2.01 \pm 0.04) \times 10^{-4}$ |
| | 3546-S-2 | $(3.43 \pm 0.04) \times 10^{-10}$ | $(1.24 \pm 0.01) \times 10^6$ | $(4.25 \pm 0.06) \times 10^{-4}$ |
| | 3548-S-1 | $(5.04 \pm 1.24) \times 10^{-10}$ | $(8.34 \pm 1.93) \times 10^5$ | $(4.05 \pm 0.19) \times 10^{-4}$ |
| | 3548-S-2 | $(3.77 \pm 2.98) \times 10^{-8}$ | $(5.31 \pm 2.15) \times 10^5$ | $(1.88 \pm 1.59) \times 10^{-2}$ |
| | 3548-S-3 | $(5.12 \pm 0.47) \times 10^{-10}$ | $(9.87 \pm 0.10) \times 10^5$ | $(5.05 \pm 0.11) \times 10^{-4}$ |
| | 3548-S-4 | $(3.26 \pm 0.93) \times 10^{-10}$ | $(2.64 \pm 0.37) \times 10^6$ | $(8.47 \pm 2.19) \times 10^{-4}$ |
| | 3548-S-5 | $(3.94 \pm 1.03) \times 10^{-10}$ | $(1.29 \pm 0.20) \times 10^6$ | $(4.99 \pm 1.08) \times 10^{-4}$ |
| | 3548-S-7 | $(2.93 \pm 1.32) \times 10^{-10}$ | $(1.58 \pm 0.28) \times 10^6$ | $(4.43 \pm 1.33) \times 10^{-4}$ |
| | 3548-S-M3-5 | $(8.71 \pm 1.98) \times 10^{-11}$ | $(2.21 \pm 1.95) \times 10^7$ | $(1.72 \pm 1.26) \times 10^{-3}$ |
| Prefusion-stabilized S (B.1.351) | 3546-S-1 | $(5.68 \pm 4.30) \times 10^{-10}$ | $(9.70 \pm 4.55) \times 10^5$ | $(4.20 \pm 0.47) \times 10^{-4}$ |
| | 3546-S-2 | $(5.40 \pm 0.03) \times 10^{-10}$ | $(1.03 \pm 0.20) \times 10^6$ | $(5.26 \pm 2.22) \times 10^{-4}$ |
| | 3548-S-1 | $(5.04 \pm 0.05) \times 10^{-10}$ | $(1.02 \pm 0.01) \times 10^6$ | $(5.13 \pm 0.11) \times 10^{-4}$ |
| | 3548-S-2 | $(1.46 \pm 1.13) \times 10^{-8}$ | $(9.82 \pm 8.35) \times 10^5$ | $(8.35 \pm 1.80) \times 10^{-3}$ |
| | 3548-S-3 | $(9.82 \pm 8.20) \times 10^{-10}$ | $(8.08 \pm 3.99) \times 10^5$ | $(5.87 \pm 2.25) \times 10^{-4}$ |
| | 3548-S-4 | $(1.05 \pm 0.23) \times 10^{-9}$ | $(9.42 \pm 1.86) \times 10^5$ | $(9.62 \pm 0.90) \times 10^{-4}$ |
| | 3548-S-5 | $(4.91 \pm 0.67) \times 10^{-10}$ | $(2.08 \pm 0.38) \times 10^6$ | $(1.02 \pm 0.21) \times 10^{-3}$ |
| | 3548-S-7 | $(2.33 \pm 0.01) \times 10^{-10}$ | $(7.64 \pm 0.03) \times 10^5$ | $(1.78 \pm 0.04) \times 10^{-4}$ |
| | 3548-S-M3-5 | $(1.23 \pm 0.99) \times 10^{-10}$ | $(2.40 \pm 1.72) \times 10^7$ | $(1.90 \pm 0.38) \times 10^{-3}$ |
| RBD (Wu) | 3546-S-2 | $(2.14 \pm 0.86) \times 10^{-10}$ | $(1.05 \pm 0.21) \times 10^6$ | $(2.13 \pm 0.56) \times 10^{-4}$ |
| | 3548-S-2 | $(6.32 \pm 1.19) \times 10^{-8}$ | $(2.22 \pm 0.17) \times 10^5$ | $(1.41 \pm 0.35) \times 10^{-2}$ |
| | 3548-RBD-15 | $(2.34 \pm 1.56) \times 10^{-8}$ | $(5.38 \pm 2.84) \times 10^4$ | $(9.77 \pm 5.15) \times 10^{-4}$ |
| RBD (B.1.351) | 3546-S-2 | $(1.59 \pm 0.68) \times 10^{-10}$ | $(3.09 \pm 0.53) \times 10^6$ | $(4.68 \pm 1.32) \times 10^{-4}$ |
| | 3548-S-2 | $(2.87 \pm 0.93) \times 10^{-8}$ | $(4.45 \pm 2.28) \times 10^5$ | $(1.14 \pm 0.25) \times 10^{-2}$ |
| HKU1 S | 3548-S-1 | $(1.26 \pm 0.07) \times 10^{-10}$ | $(6.32 \pm 0.34) \times 10^6$ | $(7.93 \pm 0.24) \times 10^{-4}$ |

## References

1. P. Zhou et al., A pneumonia outbreak associated with a new coronavirus of probable bat origin. Nature 579, 270–273 (2020).

2. M. Hoffmann et al., SARS-CoV-2 Cell Entry Depends on ACE2 and TMPRSS2 and Is Blocked by a Clinically Proven Protease Inhibitor. Cell 181, 271–280.e278 (2020).

3. Y. Cai et al., Distinct conformational states of SARS-CoV-2 spike protein. Science. 369, 1586–1592 (2020).

4. X. Fan, D. Cao, L. Kong, X. Zhang, Cryo-EM analysis of the post-fusion structure of the SARS-CoV spike glycoprotein. Nature Communications 11 (2020).

5. S. M. Costello et al., The SARS-CoV-2 spike reversibly samples an open-trimer conformation exposing novel epitopes. Nature Structural & Molecular Biology 29, 229–238 (2022).

6. F. P. Polack et al., Safety and Efficacy of the BNT162b2 mRNA Covid-19 Vaccine. New England Journal of Medicine 383, 2603–2615 (2020).

7. L. R. Baden et al., Efficacy and Safety of the mRNA-1273 SARS-CoV-2 Vaccine. New England Journal of Medicine 384, 403–416 (2021).

8. J. Sadoff et al., Safety and Efficacy of Single-Dose Ad26.COV2.S Vaccine against Covid-19. New England Journal of Medicine 384, 2187–2201 (2021).

9. J. Pallesen et al., Immunogenicity and structures of a rationally designed prefusion MERS-CoV spike antigen. Proceedings of the National Academy of Sciences 114, E7348–E7357 (2017).

10. S. Yang et al., Safety and immunogenicity of a recombinant tandem-repeat dimeric RBD-based protein subunit vaccine (ZF2001) against COVID-19 in adults: two randomised, double-blind, placebo-controlled, phase 1 and 2 trials. The Lancet Infectious Diseases 10.1016/s1473-3099(21)00127-4 (2021).

11. M. Pino et al., A yeast expressed RBD-based SARS-CoV-2 vaccine formulated with 3M-052-alum adjuvant promotes protective efficacy in non-human primates. Science Immunology 6, eabh3634 (2021).

12. N. C. Kyriakidis, A. López-Cortés, E. V. González, A. B. Grimaldos, E. O. Prado, SARS-CoV-2 vaccines strategies: a comprehensive review of phase 3 candidates. npj Vaccines 6 (2021).

13. Y. Liao et al., Safety and immunogenicity of a recombinant interferon-armed RBD dimer vaccine (V-01) for COVID-19 in healthy adults: a randomized, double-blind, placebo-controlled, Phase I trial. Emerging Microbes & Infections 10, 1589–1597 (2021).

14. K. Xu et al., Recombinant chimpanzee adenovirus AdC7 expressing dimeric tandem-repeat spike protein RBD protects mice against COVID-19. Emerging Microbes & Infections 10, 1574–1588 (2021).

15. J. Pollet et al., Receptor-binding domain recombinant protein on alum-CpG induces broad protection against SARS-CoV-2 variants of concern. Vaccine. 40, 3655–3663 (2022).

16. B. Ju et al., Human neutralizing antibodies elicited by SARS-CoV-2 infection. Nature 584, 115–119 (2020).

17. P. J. M. Brouwer et al., Potent neutralizing antibodies from COVID-19 patients define multiple targets of vulnerability. Science 369, 643–650 (2020).

18. A. Deshpande, B. D. Harris, L. Martinez-Sobrido, J. J. Kobie, M. R. Walter, Epitope Classification and RBD Binding Properties of Neutralizing Antibodies Against SARS-CoV-2 Variants of Concern. Frontiers in immunology. 12 (2021).

19. A. Abayasingam et al., Long-term persistence of RBD+ memory B cells encoding neutralizing antibodies in SARS-CoV-2 infection. Cell Reports Medicine 2, 100228 (2021).

20. A. Sokal et al., Maturation and persistence of the anti-SARS-CoV-2 memory B cell response. Cell 184, 1201–1213 e1214 (2021).

21. Z. Wang et al., Naturally enhanced neutralizing breadth against SARS-CoV-2 one year after infection. Nature 595, 426–431 (2021).

22. P. Kotagiri et al., B cell receptor repertoire kinetics after SARS-CoV-2 infection and vaccination. Cell Reports 38, 110393 (2022).

23. M. S. A. Gilman et al., Rapid profiling of RSV antibody repertoires from the memory B cells of naturally infected adult donors. Science Immunology 1, eaaj1879–eaaj1818 (2016).

24. P. Tong et al., Memory B cell repertoire for recognition of evolving SARS-CoV-2 spike. Cell 184, 4969–4980.e4915 (2021).

25. A. Facciuolo et al., High-resolution analysis of long-term serum antibodies in humans following convalescence of SARS-CoV-2 infection. Scientific Reports 12 (2022).

26. W. N. Voss et al., Prevalent, protective, and convergent IgG recognition of SARS-CoV-2 non-RBD spike epitopes. Science 372, 1108–1112 (2021).

27. B. Isho et al., Persistence of serum and saliva antibody responses to SARS-CoV-2 spike antigens in patients with COVID-19. Science Immunology 5, eabe5511 (2020).

28. A. S. Iyer et al., Persistence and decay of human antibody responses to the receptor binding domain of SARS-CoV-2 spike protein in COVID-19 patients. Science Immunology 5, eabe0367 (2020).

29. A. Wajnberg et al., Robust neutralizing antibodies to SARS-CoV-2 infection persist for months. Science 370, 1227–1230 (2020).

30. A. Haveri et al., Persistence of neutralizing antibodies a year after SARS-CoV-2 infection in humans. European Journal of Immunology 51, 3202–3213 (2021).

31. E. M. Anderson et al., Seasonal human coronavirus antibodies are boosted upon SARS-CoV-2 infection but not associated with protection. Cell 184, 1858–1864.e1810 (2021).

32. P. R. Wratil et al., Evidence for increased SARS-CoV-2 susceptibility and COVID-19 severity related to pre-existing immunity to seasonal coronaviruses. Cell Reports 37, 110169 (2021).

33. T. Woudenberg et al., Humoral immunity to SARS-CoV-2 and seasonal coronaviruses in children and adults in north-eastern France. EBioMedicine. 70, 103495 (2021).

34. K. W. Ng et al., Preexisting and de novo humoral immunity to SARS-CoV-2 in humans. Science. 370, 1339–1343 (2020).

35. P. Nguyen-Contant et al., S Protein-Reactive IgG and Memory B Cell Production after Human SARS-CoV-2 Infection Includes Broad Reactivity to the S2 Subunit. MBio. 11 (2020).

36. T. Aydillo et al., Immunological imprinting of the antibody response in COVID-19 patients. Nature Communications 12 (2021).

37. M. Aguilar-Bretones et al., Seasonal coronavirus-specific B-cells with limited SARS-CoV-2 cross-reactivity dominate the IgG response in severe COVID-19 patients. Journal of Clinical Investigation 10.1172/jci150613 (2021).

38. M. Sakharkar et al., Prolonged evolution of the human B cell response to SARS-CoV-2 infection. Science Immunology 6, eabg6916 (2021).

39. A. Z. Wec et al., Broad neutralization of SARS-related viruses by human monoclonal antibodies. Science 369, 731–736 (2020).

40. T. N. Starr et al., Prospective mapping of viral mutations that escape antibodies used to treat COVID-19. Science 371, 850–854 (2021).

41. J. A. Plante et al., Spike mutation D614G alters SARS-CoV-2 fitness. Nature 592, 116–121 (2021).

42. H. Liu et al., The basis of a more contagious 501Y.V1 variant of SARS-CoV-2. Cell Research 31, 720–722 (2021).

43. B. Korber et al., Tracking Changes in SARS-CoV-2 Spike: Evidence that D614G Increases Infectivity of the COVID-19 Virus. Cell 182, 812–827.e819 (2020).

44. A. Khan et al., Higher infectivity of the SARS-CoV-2 new variants is associated with K417N/T, E484K, and N501Y mutants: An insight from structural data. Journal of Cellular Physiology 10.1002/jcp.30367 (2021).

45. Y. J. Hou et al., SARS-CoV-2 D614G variant exhibits efficient replication ex vivo and transmission in vivo. Science 10.1126/science.abe8499, eabe8499 (2020).

46. Q. Li et al., The Impact of Mutations in SARS-CoV-2 Spike on Viral Infectivity and Antigenicity. Cell 182, 1284–1294.e1289 (2020).

47. X. Zhang et al., SARS-CoV-2 Omicron strain exhibits potent capabilities for immune evasion and viral entrance. Signal Transduction and Targeted Therapy 6 (2021).

48. V. V. Edara et al., Infection- and vaccine-induced antibody binding and neutralization of the B.1.351 SARS-CoV-2 variant. Cell Host & Microbe 29, 516–521.e513 (2021).

49. A. J. Greaney et al., Comprehensive mapping of mutations in the SARS-CoV-2 receptor-binding domain that affect recognition by polyclonal human plasma antibodies. Cell Host & Microbe 29, 463–476.e466 (2021).

50. M. Yuan et al., Structural and functional ramifications of antigenic drift in recent SARS-CoV-2 variants. Science 10.1126/science.abh1139, eabh1139 (2021).

51. M. Widera et al., Limited neutralization of authentic SARS-CoV-2 variants carrying E484K in vitro. The Journal of Infectious Diseases 10.1093/infdis/jiab355 (2021).

52. C. K. Wibmer et al., SARS-CoV-2 501Y.V2 escapes neutralization by South African COVID-19 donor plasma. Nature Medicine 27, 622–625 (2021).

53. P. Wang et al., Antibody resistance of SARS-CoV-2 variants B.1.351 and B.1.1.7. Nature 593, 130–135 (2021).

54. M. Becker et al., Immune response to SARS-CoV-2 variants of concern in vaccinated individuals. Nature Communications 12 (2021).

55. T. N. Starr, A. J. Greaney, A. S. Dingens, J. D. Bloom, Complete map of SARS-CoV-2 RBD mutations that escape the monoclonal antibody LY-CoV555 and its cocktail with LY-CoV016. Cell Reports Medicine 2, 100255 (2021).

56. L. Zhang et al., SARS-CoV-2 spike-protein D614G mutation increases virion spike density and infectivity. Nature Communications 11 (2020).

57. H. Liu et al., 501Y.V2 and 501Y.V3 variants of SARS-CoV-2 lose binding to bamlanivimab in vitro. mAbs 13, 1919285 (2021).

58. R. E. Chen et al., Resistance of SARS-CoV-2 variants to neutralization by monoclonal and serum-derived polyclonal antibodies. Nature Medicine 27, 717–726 (2021).

59. Z. Liu et al., Identification of SARS-CoV-2 spike mutations that attenuate monoclonal and serum antibody neutralization. Cell Host & Microbe 29, 477–488.e474 (2021).

60. F. J. Ibarrondo et al., Rapid Decay of Anti–SARS-CoV-2 Antibodies in Persons with Mild Covid-19. New England Journal of Medicine 383, 1085–1087 (2020).

61. B. Zhou et al., SARS-CoV-2 spike D614G change enhances replication and transmission. Nature 592, 122–127 (2021).

62. X. Deng et al., Transmission, infectivity, and neutralization of a spike L452R SARS-CoV-2 variant. Cell 184, 3426–3437.e3428 (2021).

63. B. Li et al., Viral infection and transmission in a large, well-traced outbreak caused by the SARS-CoV-2 Delta variant. Nature Communications 13 (2022).

64. D. Planas et al., Sensitivity of infectious SARS-CoV-2 B.1.1.7 and B.1.351 variants to neutralizing antibodies. Nature Medicine 27, 917–924 (2021).

65. P. Arora et al., Delta variant (B.1.617.2) sublineages do not show increased neutralization resistance. Cellular & Molecular Immunology 18, 2557–2559 (2021).

66. D. Planas et al., Reduced sensitivity of SARS-CoV-2 variant Delta to antibody neutralization. Nature 596, 276–280 (2021).

67. L. Cheng et al., Cross-neutralization of SARS-CoV-2 Kappa and Delta variants by inactivated vaccine-elicited serum and monoclonal antibodies. Cell Discovery 7 (2021).

68. D. Planas et al., Considerable escape of SARS-CoV-2 Omicron to antibody neutralization. Nature 10.1038/s41586-021-04389-z (2021).

69. L. A. Vanblargan et al., An infectious SARS-CoV-2 B.1.1.529 Omicron virus escapes neutralization by therapeutic monoclonal antibodies. Nature Medicine 10.1038/s41591-021-01678-y (2022).

70. A. Choi et al., Safety and immunogenicity of SARS-CoV-2 variant mRNA vaccine boosters in healthy adults: an interim analysis. Nature Medicine 27, 2025–2031 (2021).

71. S. Chiba et al., Multivalent nanoparticle-based vaccines protect hamsters against SARS-CoV-2 after a single immunization. Communications Biology 4 (2021).

72. S. A. Kim et al., A Multivalent Vaccine Based on Ferritin Nanocage Elicits Potent Protective Immune Responses against SARS-CoV-2 Mutations. International Journal of Molecular Sciences 23, 6123 (2022).

73. C. Sun et al., Induction of Broadly Cross-Reactive Antibody Responses to SARS-CoV-2 Variants by S1 Nanoparticle Vaccines. Journal of virology 96, e0038322–e0038322 (2022).

74. K. W. Ng et al., SARS-CoV-2 S2-targeted vaccination elicits broadly neutralizing antibodies. Science translational medicine 14, 1–eabn3715 (2022).

75. P. J. Halfmann, et al., Potent neutralization of SARS-CoV-2 including variants of concern by vaccines presenting the receptor-binding domain multivalently from nanoscaffolds. Bioengineering & translational medicine 6, e10253-n/a (2021).

76. B. Aparicio et al., Preclinical evaluation of a synthetic peptide vaccine against SARS-CoV-2 inducing multiepitopic and cross-reactive humoral neutralizing and cellular CD4 and CD8 responses. Emerging microbes & infections 10, 1931–1946 (2021).

77. D. R. Boutz et al., Proteomic Identification of Monoclonal Antibodies from Serum. Analytical Chemistry 86, 4758–4766 (2014).

78. J. J. Lavinder et al., Identification and characterization of the constituent human serum antibodies elicited by vaccination. Proceedings of the National Academy of Sciences 111, 2259–2264 (2014).

79. J. Lee et al., Molecular-level analysis of the serum antibody repertoire in young adults before and after seasonal influenza vaccination. Nature Medicine 22, 1456–1464 (2016).

80. J. Lee et al., Persistent Antibody Clonotypes Dominate the Serum Response to Influenza over Multiple Years and Repeated Vaccinations. Cell Host & Microbe 25, 367–376.e365 (2019).

81. A. R. Crowley et al., Boosting of cross-reactive antibodies to endemic coronaviruses by SARS-CoV-2 infection but not vaccination with stabilized spike. eLife 11 (2022).

82. H. Natarajan et al., Markers of Polyfunctional SARS-CoV-2 Antibodies in Convalescent Plasma. MBio. 12 (2021).

83. S. E. Butler et al., Distinct Features and Functions of Systemic and Mucosal Humoral Immunity Among SARS-CoV-2 Convalescent Individuals. Front Immunol 11, 618685 (2020).

84. C. L. Hsieh et al., Structure-based design of prefusion-stabilized SARS-CoV-2 spikes. Science 369, 1501–1505 (2020).

85. L. Liu et al., Potent neutralizing antibodies against multiple epitopes on SARS-CoV-2 spike. Nature 584, 450–456 (2020).

86. R. D. Melani et al., Next-Generation Serology by Mass Spectrometry: Readout of the SARS-CoV-2 Antibody Repertoire. Journal of Proteome Research 21, 274–288 (2022).

87. J. Jung et al., Influenza vaccination in the elderly boosts antibodies against conserved viral proteins and egg-produced glycans. Journal of Clinical Investigation 131 (2021).

88. M. Kuraoka et al., Infant Antibody Repertoires during the First Two Years of Influenza Vaccination. MBio. 13 (2022).

89. C. Gaebler et al., Evolution of antibody immunity to SARS-CoV-2. Nature 591, 639–644 (2021).

90. Y. Chen et al., Quick COVID-19 Healers Sustain Anti-SARS-CoV-2 Antibody Production. Cell. 183, 1496–1507.e1416 (2020).

91. T. Hattori et al., The ACE2-binding Interface of SARS-CoV-2 Spike Inherently Deflects Immune Recognition. J Mol Biol 433, 166748 (2021).

92. H. Ma et al., Potent Neutralization of SARS-CoV-2 by Hetero-Bivalent Alpaca Nanobodies Targeting the Spike Receptor-Binding Domain. Journal of virology : 95 (2021).

93. A. J. Greaney et al., Antibodies elicited by mRNA-1273 vaccination bind more broadly to the receptor binding domain than do those from SARS-CoV-2 infection. Science translational medicine 13 (2021).

94. K. Nayak et al., Characterization of neutralizing versus binding antibodies and memory B cells in COVID-19 recovered individuals from India. *Virology (New York*, N.Y.) 558, 13–21 (2021).

95. S. C. A. Nielsen et al., Human B Cell Clonal Expansion and Convergent Antibody Responses to SARS-CoV-2. Cell Host & Microbe 28, 516–525.e515 (2020).

96. J. M. Dan et al., Immunological memory to SARS-CoV-2 assessed for up to 8 months after infection. Science 371, eabf4063 (2021).

97. X.-L. Jiang et al., Lasting antibody and T cell responses to SARS-CoV-2 in COVID-19 patients three months after infection. Nature Communications 12 (2021).

98. A. K. Wheatley et al., Evolution of immune responses to SARS-CoV-2 in mild-moderate COVID-19. Nature Communications 12 (2021).

99. J. S. Turner et al., SARS-CoV-2 infection induces long-lived bone marrow plasma cells in humans. Nature 595, 421–425 (2021).

100. W. E. Harrington et al., Rapid decline of neutralizing antibodies is associated with decay of IgM in adults recovered from mild COVID-19. Cell Reports Medicine 2, 100253 (2021).

101. S. Yoshida et al., SARS-CoV-2-induced humoral immunity through B cell epitope analysis in COVID-19 infected individuals. Scientific Reports 11 (2021).

102. B. Bošnjak et al., Low serum neutralizing anti-SARS-CoV-2 S antibody levels in mildly affected COVID-19 convalescent patients revealed by two different detection methods. Cellular & Molecular Immunology 18, 936–944 (2021).

103. C. Rydyznski Moderbacher et al., Antigen-Specific Adaptive Immunity to SARS-CoV-2 in Acute COVID-19 and Associations with Age and Disease Severity. Cell 183, 996–1012.e1019 (2020).

104. E. Salazar et al., Convalescent plasma anti–SARS-CoV-2 spike protein ectodomain and receptor-binding domain IgG correlate with virus neutralization. Journal of Clinical Investigation 130, 6728–6738 (2020).

105. L. Ni et al., Detection of SARS-CoV-2-Specific Humoral and Cellular Immunity in COVID-19 Convalescent Individuals. Immunity 52, 971–977.e973 (2020).

106. D. Sterlin et al., IgA dominates the early neutralizing antibody response to SARS-CoV-2. Sci Transl Med 13 (2021).

107. R. Gasser et al., Major role of IgM in the neutralizing activity of convalescent plasma against SARS-CoV-2. Cell Reports 34, 108790 (2021).

108. J. Shang et al., Cell entry mechanisms of SARS-CoV-2. Proceedings of the National Academy of Sciences 117, 11727–11734 (2020).

109. X. Huang et al., Human Coronavirus HKU1 Spike Protein UsesO-Acetylated Sialic Acid as an Attachment Receptor Determinant and Employs Hemagglutinin-Esterase Protein as a Receptor-Destroying Enzyme. Journal of Virology 89, 7202–7213 (2015).

110. J. Hicks et al., Serologic Cross-Reactivity of SARS-CoV-2 with Endemic and Seasonal Betacoronaviruses. Journal of Clinical Immunology 10.1007/s10875-021-00997-6 (2021).

111. D. Li et al., In vitro and in vivo functions of SARS-CoV-2 infection-enhancing and neutralizing antibodies. Cell 184, 4203–4219 e4232 (2021).

112. Y. Liu et al., An infectivity-enhancing site on the SARS-CoV-2 spike protein targeted by antibodies. Cell 184, 3452–3466 e3418 (2021).

113. Y. Wu et al., Identification of Human Single-Domain Antibodies against SARS-CoV-2. Cell Host & Microbe 27, 891–898.e895 (2020).

114. F. Lucca et al., Immunogenicity and Safety of the BNT162b2 COVID-19 Vaccine in Patients with Cystic Fibrosis with or without Lung Transplantation. International Journal of Molecular Sciences 24, 908 (2023).

115. G. Alicandro et al., Immunogenicity of BNT162b2 mRNA-Based Vaccine against SARS-CoV-2 in People with Cystic Fibrosis According to Disease Characteristics and Maintenance Therapies. Biomedicines 10, 1998 (2022).

116. A. Michos et al., Immunogenicity of the COVID-19 BNT162b2 vaccine in adolescents and young adults with cystic fibrosis. Journal of Cystic Fibrosis 21, e184–e187 (2022).

117. M. Hoffmann et al., Profound neutralization evasion and augmented host cell entry are hallmarks of the fast-spreading SARS-CoV-2 lineage XBB.1.5. Cellular & Molecular Immunology 10.1038/s41423-023-00988-0 (2023).

118. M. J. Makela et al., Viruses and bacteria in the etiology of the common cold. Journal of clinical microbiology 36, 539–542 (1998).

119. R. Dijkman et al., The dominance of human coronavirus OC43 and NL63 infections in infants. Journal of clinical virology : the official publication of the Pan American Society for Clinical Virology 53, 135–139 (2012).

120. W. Zhou, W. Wang, H. Wang, R. Lu, W. Tan, First infection by all four non-severe acute respiratory syndrome human coronaviruses takes place during childhood. BMC infectious diseases 13, 433 (2013).

121. K. Tamminen, M. Salminen, V. Blazevic, Seroprevalence and SARS-CoV-2 cross-reactivity of endemic coronavirus OC43 and 229E antibodies in Finnish children and adults. Clinical immunology. 229, 108782 (2021).

122. J. T. Sandberg et al., SARS-CoV-2-specific humoral and cellular immunity persists through 9 months irrespective of COVID-19 severity at hospitalisation. Clinical & Translational Immunology 10 (2021).

123. G. C. Ippolito et al., Antibody Repertoires in Humanized NOD-scid-IL2Rγnull Mice and Human B Cells Reveals Human-Like Diversification and Tolerance Checkpoints in the Mouse. PLoS ONE 7, e35497 (2012).

124. M. Letko, A. Marzi, V. Munster, Functional assessment of cell entry and receptor usage for SARS-CoV-2 and other lineage B betacoronaviruses. Nat Microbiol 5, 562–569 (2020).

125. M. E. Ackerman et al., A robust, high-throughput assay to determine the phagocytic activity of clinical antibody samples. Journal of Immunological Methods 366, 8–19 (2011).

126. E. G. McAndrew et al., Determining the Phagocytic Activity of Clinical Antibody Samples. Journal of Visualized Experiments 10.3791/3588 (2011).

127. S. H. Tam et al., Functional, Biophysical, and Structural Characterization of Human IgG1 and IgG4 Fc Variants with Ablated Immune Functionality. Antibodies (Basel) 6 (2017).

128. S. Fischinger et al., A high-throughput, bead-based, antigen-specific assay to assess the ability of antibodies to induce complement activation. Journal of immunological methods. 473, 112630 (2019).

